# Intrinsic functional connectivity of the central extended amygdala

**DOI:** 10.1101/178533

**Authors:** Rachael M. Tillman, Melissa D. Stockbridge, Brendon M. Nacewicz, Salvatore Torrisi, Andrew S. Fox, Jason F. Smith, Alexander J. Shackman

**Affiliations:** Department of Psychology, University of Maryland, College Park, MD 20742 USA.; Department of Hearing and Speech Sciences, University of Maryland, College Park, MD 20742 USA.; Neuroscience and Cognitive Science Program, University of Maryland, College Park, MD 20742 USA.; Maryland Neuroimaging Center, University of Maryland, College Park, MD 20742 USA.; Department of Psychiatry, University of Wisconsin—Madison, 6001 Research Park Boulevard, Madison, WI 53719 USA.; Section on the Neurobiology of Fear and Anxiety, National Institute of Mental Health, Bethesda, MD 20892 USA.; Department of Psychology, University of California, Davis, CA 95616 USA.; California National Primate Research Center, University of California, Davis, CA 95616 USA.

**Keywords:** affective neuroscience, bed nucleus of the stria terminalis (BST), central extended amygdala, Nathan Kline Institute-Rockland Sample (NKI-RS), resting-state fMRI

## Abstract

The central extended amygdala (EAc)—including the bed nucleus of the stria terminalis (BST) and central nucleus of the amygdala (Ce)—plays a key role in orchestrating states of fear and anxiety and is implicated in the development and maintenance of anxiety disorders, depression, and substance abuse. Although it is widely thought that these disorders reflect the coordinated actions of large-scale functional circuits in the brain, the architecture of the EAc functional network, and the degree to which the BST and the Ce show distinct patterns of intrinsic functional connectivity, remains incompletely understood. Here, we leveraged a combination of approaches to trace the connectivity of the BST and the Ce in 130 psychiatrically healthy, racially diverse, community-dwelling adults with enhanced power and precision. Multiband imaging, high-precision data registration techniques, and spatially unsmoothed data were used to maximize anatomical specificity. Using newly developed seed regions, whole-brain regression analyses revealed robust functional connectivity between the BST and Ce via the sublenticular extended amygdala (‘substantia innominata’), the ribbon of subcortical gray matter encompassing the ventral amygdalofugal pathway. Both regions displayed significant coupling with the ventromedial prefrontal cortex (vmPFC), midcingulate cortex (MCC), insula, and anterior hippocampus. The BST showed significantly stronger connectivity with prefrontal territories—including the vmPFC, anterior MCC and pregenual anterior cingulate cortex—as well as the thalamus, striatum, and the periaqueductal gray. The only regions showing stronger functional connectivity with the Ce were located in the anterior hippocampus and dorsal amygdala. These observations provide a baseline against which to compare a range of special populations, inform our understanding of the role of the EAc in normal and pathological fear and anxiety, and highlight the value of several new approaches to image registration which may be particularly useful for researchers working with ‘de-identified’ neuroimaging data.

**GRAPHICAL ABSTRACT:** Intrinsic functional connectivity of bed nucleus of the stria terminalis (BST) and the central nucleus of the amygdala (Ce) in 130 psychiatrically healthy adults.

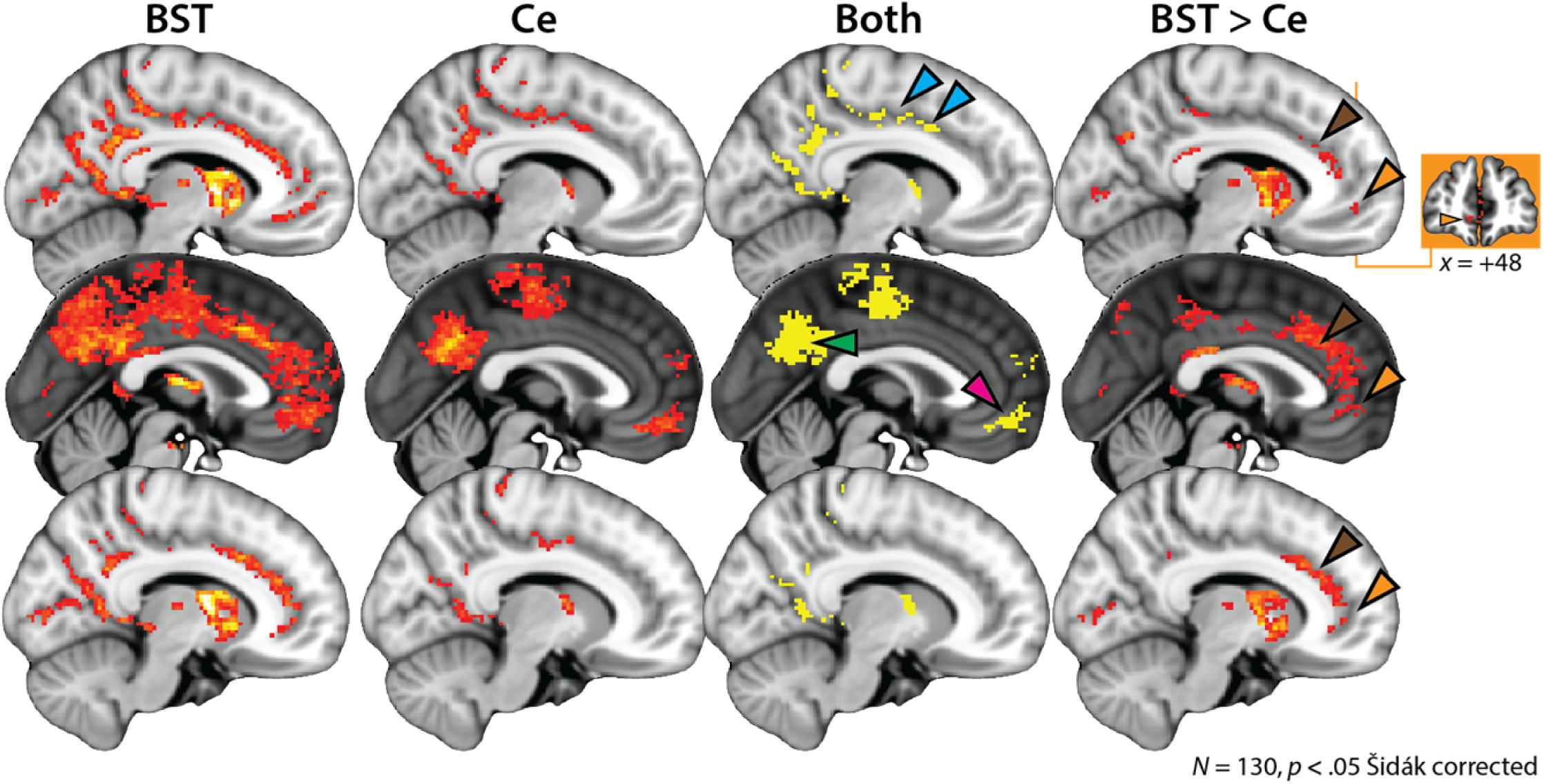

**HIGHLIGHTS:**

- BST and Ce implicated in normal and pathological fear and anxiety
- Traced the intrinsic functional connectivity of the BST and the Ce in 130 adults
- Multiband imaging, high-precision registration, unsmoothed data, newly developed seeds
- BST and Ce show robust coupling with one another, hippocampus, insula, MCC, and vmPFC
- BST shows stronger coupling with prefrontal/cingulate territories and brainstem/PAG

## INTRODUCTION

When extreme, fear and anxiety can become debilitating (Grupe & Nitschke, 2013; Salomon et al., 2015). Anxiety disorders are common and challenging to treat, imposing a staggering burden on public health, and underscoring the need to develop a deeper understanding of the distributed neural circuits governing the expression of fear and anxiety in humans (Bystritsky, 2006; DiLuca & Olesen, 2014; Griebel & Holmes, 2013; Wang, Gaitsch, Poon, Cox, & Rzhetsky, *in press;* Whiteford et al., 2013).

Converging lines of anatomical, mechanistic, and physiological evidence make it clear that the central extended amygdala (EAc) is a key hub in this circuitry (**Figure 1a,b**) (Avery, Clauss, & Blackford, 2016; Davis, Walker, Miles, & Grillon, 2010; A. S. Fox & Shackman, *under review;* Goode & Maren, 2017; Gungor & Paré, 2016; Shackman & Fox, 2016; Tovote, Fadok, & Luthi, 2015). The EAc encompasses a collection of subcortical regions with similar cellular compositions, neurochemistry, gene expression, and structural connectivity and encompasses the bed nucleus of the stria terminalis (BST), the central nucleus of the amygdala (Ce), the sublenticular extended amygdala (SLEA), and portions of the accumbens shell (Alheid & Heimer, 1988; A. S. Fox, Oler, Tromp, Fudge, & Kalin, 2015; Oler et al., 2017; Yilmazer-Hanke, 2012). It has long been recognized that the amygdala is connected to the BST via two major fiber bundles—the ventral amygdalofugal pathway (VA) and the stria terminalis (ST) (Avery et al., 2014; Kamali et al., 2016; Kamali et al., 2015; Nauta, 1961) (**Figure 1c**)—and more recent tracing studies have identified a third, indirect pathway centered on the SLEA (Ce ↔ SLEA ↔ BSTL) (deCampo & Fudge, 2013; Oler et al., 2017). Anatomically, the Ce and the BST are both poised to trigger or orchestrate key signs of fear and anxiety— including alterations in arousal, behavioral inhibition, and neuroendocrine activity—via dense mono- and poly-synaptic projections to brainstem and subcortical effector regions (A. S. Fox, Oler, Tromp, et al., 2015; Freese & Amaral, 2009).

**Figure 1.**
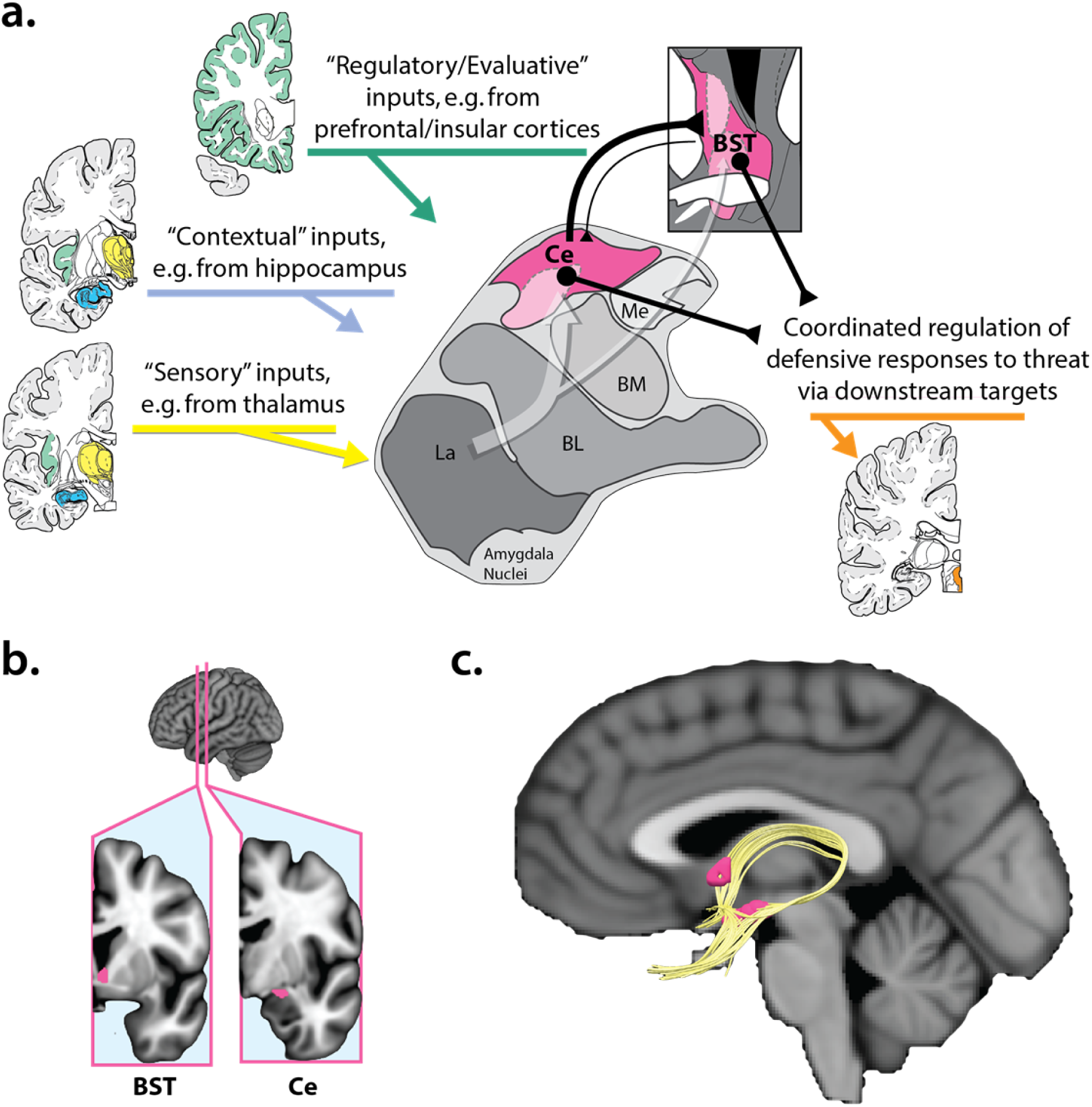
The EAc. a. Simplified schematic of key EAc inputs and outputs in humans and other primates. The EAc (*magenta*) encompasses the BST, which encircles the anterior commissure, and the Ce. As shown by the translucent white arrow at the center of the figure, much of the sensory (*yellow*), contextual (*blue*), and regulatory (*green*) inputs to the EAc are indirect (i.e., polysynaptic), and often first pass through adjacent amygdala nuclei before arriving at the Ce or the BST. Both regions are poised to orchestrate momentary states of fear and anxiety via dense projections to downstream effector regions (*orange*). Portions of this figure were adapted from the atlas of (Mai, Paxinos, & Voss, 2007; see also Yilmazer-Hanke, 2012). ***b. BST and Ce seeds.*** Figure depicts the location of the BST and Ce seeds used in the present study. See **Supplementary Figure S5** for bilateral views and a more detailed description of seed derivation. ***c. Structural connections of the EAc.*** In humans and other primates, the BST (*dorsorostral magenta region*) and the Ce (*ventrocaudal magenta region*) are structurally connected via two major fiber bundles (*gold*), the ventral amygdalofugal pathway and the stria terminalis (Johnston, 1923; Nauta, 1961; Yilmazer-Hanke, 2012). From the Ce, the ventral amygdalofugal pathway courses forward and medially, passing through the SLEA, a bridge of neurons harbored within the substantia innominata. The stria terminalis, which arches dorsally over the thalamus, provides a second, less direct connection between the two major divisions of the central extended amygdala. Figure depicts deterministic tractography (*gold*) of these two fiber bundles. Image kindly provided by Do Tromp. Abbreviations—BL, basolateral nucleus of the amygdala; BM, basomedial nucleus of the amygdala; BST, bed nucleus of the stria terminalis; Ce, central nucleus of the amygdala; EAc, central division of the extended amygdala; La, lateral nucleus of the amygdala; Me, medial nucleus of the amygdala; SLEA, sublenticular extended amygdala.

Consistent with this neuroanatomy, mechanistic studies in rodents indicate that microcircuits within and between the BST and the Ce play a critical role in organizing defensive responses to a range of potentially threat-relevant cues and contexts (Calhoon & Tye, 2015; Davis et al., 2010; A. S. Fox & Shackman, *under review;* Goode & Maren, 2017; Gungor & Paré, 2016; Lange et al., *in press;* Tovote et al., 2015) (**Figure 1c**). Although the BST and the Ce are often viewed as passive relays for amygdala-mediated emotional learning (e.g., L → Ce/BST → effector regions; LeDoux, 2000, 2007; Pare & Duvarci, 2012), more recent work in rodents has expanded this role to include guiding attention to motivationally salient stimuli (Davis & Whalen, 2001; Roesch, Esber, Li, Daw, & Schoenbaum, 2012; Shackman, Kaplan, et al., 2016), learning aversive associations (Ciocchi et al., 2010; Han, Soleiman, Soden, Zweifel, & Palmiter, 2015; Li et al., 2013; Penzo, Robert, & Li, 2014; Penzo et al., 2015; Sato et al., 2015; Yu et al., 2017), and actively gating and regulating defensive responses (Ehrlich et al., 2009; Fadok et al., 2017; Gungor & Paré, 2016; Pare & Duvarci, 2012).

Although the causal contribution of the BST has yet to be explored in primates, the Ce has been shown to control defensive responses to potential threat in monkeys (Kalin, 2017; Kalin et al., 2016; Kalin, Shelton, & Davidson, 2004). Likewise, rodents, monkeys, and humans with amygdala damage exhibit a profound lack of fear and anxiety in response to a broad spectrum of learned and innate dangers (Antoniadis, Winslow, Davis, & Amaral, 2007; Bechara et al., 1995; J. S. Choi & Kim, 2010; Davis & Whalen, 2001; Feinstein, Adolphs, Damasio, & Tranel, 2011; Feinstein, Adolphs, & Tranel, 2016; Izquierdo, Suda, & Murray, 2005; Kalin et al., 2004; Korn et al., *in press;* Mason, Capitanio, Machado, Mendoza, & Amaral, 2006; Oler, Fox, Shackman, & Kalin, 2016).

Neuroimaging research indicates that heightened activity in the EAc is associated with elevated signs of fear and anxiety in both monkeys and humans (Alvarez et al., 2015; Banihashemi, Sheu, Midei, & Gianaros, 2015; Cheng, Knight, Smith, & Helmstetter, 2006; Cheng, Richards, & Helmstetter, 2007; A. S. Fox, Oler, Shackman, et al., 2015; A. S. Fox, Shelton, Oakes, Davidson, & Kalin, 2008; Kalin, Shelton, Fox, Oakes, & Davidson, 2005; Knight, Nguyen, & Bandettini, 2005; Kragel & LaBar, 2015; LaBar, Gatenby, Gore, LeDoux, & Phelps, 1998; Shackman et al., 2013; Somerville et al., 2013; van Well, Visser, Scholte, & Kindt, 2012; Wood, Ver Hoef, & Knight, 2014). Among humans, the amygdala responds to a variety of threat-related cues (Costafreda, Brammer, David, & Fu, 2008; Fusar-Poli et al., 2009; Lindquist, Satpute, Wager, Weber, & Barrett, 2016; Sabatinelli et al., 2011; Sergerie, Chochol, & Armony, 2008) and work using high-resolution fMRI indicates that the dorsal amygdala in the region of the Ce is particularly sensitive to aversive visual stimuli (Hrybouski et al., 2016). Although less intensively studied than the Ce, the BST is sensitive to emotional faces (Sladky et al., *in press*), aversive images (Brinkmann et al., *under review/personal communication 7/20/2017*), and a variety of threat-related cues (Alvarez, Chen, Bodurka, Kaplan, & Grillon, 2011; Brinkmann et al., 2017; J. M. Choi, Padmala, & Pessoa, 2012; Grupe, Oathes, & Nitschke, 2013; Herrmann et al., 2016; Klumpers et al., 2015; McMenamin, Langeslag, Sirbu, Padmala, & Pessoa, 2014; Mobbs et al., 2010; Pedersen et al., 2017; Somerville et al., 2013; Somerville, Whalen, & Kelley, 2010). While imaging research hints at functional differences between the two regions (e.g., Alvarez et al., 2011; A. S. Fox, Oler, Shackman, et al., 2015; Shackman et al., 2017; Somerville et al., 2013), methodological limitations preclude decisive inferences (A. S. Fox & Shackman, *under review;* Shackman & Fox, 2016). Importantly, other work suggests that alterations in EAc function likely plays a key role in the development, maintenance, and recurrence of anxiety disorders, depression, and substance abuse (Avery et al., 2016; A. S. Fox & Kalin, 2014; Kaczkurkin et al., 2016; Shackman, Kaplan, et al., 2016; Shackman, Tromp, et al., 2016; Stevens et al., 2017; Wise & Koob, 2014).

Although this vast literature leaves little doubt that the EAc plays a crucial role in evaluating and responding to a variety of potential threats, it does not act in isolation. Fear and anxiety reflect functional circuits that extend well beyond the borders of the EAc (e.g., Chang, Gianaros, Manuck, Krishnan, & Wager, 2015; A. S. Fox & Shackman, *under review;* Kragel, Knodt, Hariri, & LaBar, 2016; Nummenmaa & Saarimaki, *in press;* Pessoa, 2017; Shackman & Fox, *in press;* Shackman, Fox, & Seminowicz, 2015; Wager et al., 2015). Anatomically, the BST and the Ce are embedded within a complex web of mono- and polysynaptically connected brain regions (**Figure 1a**) (Carrive & Morgan, 2012; A. S. Fox, Oler, Tromp, et al., 2015; Freese & Amaral, 2009; Oler et al., 2017; Ongur & Price, 2000). This structural backbone includes subcortical regions, such as the periaqueductal gray (PAG), that are responsible for triggering specific signs of fear and anxiety (Amano et al., 1982; Assareh, Sarrami, Carrive, & McNally, 2016; Bandler, Price, & Keay, 2000; Chen et al., 2015; Fadok et al., 2017; Faull & Pattinson, 2017; Motta, Carobrez, & Canteras, 2017; Nashold, Wilson, & Slaughter, 1969; Richardson & Akil, 1977; Satpute et al., 2013; Tovote et al., 2016). It also encompasses a number of cortical regions implicated in fear and anxiety, including the anterior insula, dorsolateral prefrontal cortex, mid-cingulate cortex (MCC), and OFC (e.g., Birn et al., 2014; Buhle et al., 2014; Cavanagh & Shackman, 2015; de la Vega, Chang, Banich, Wager, & Yarkoni, 2016; A. S. Fox, Oler, Shackman, et al., 2015; A. S. Fox et al., 2010; Grupe & Nitschke, 2013; Shackman, McMenamin, Maxwell, Greischar, & Davidson, 2009; Shackman et al., 2011; Stout, Shackman, Pedersen, Miskovich, & Larson, *in press;* Uddin, Kinnison, Pessoa, & Anderson, 2014). While it is widely believed that the synchronized flow of information across this network underlies the human capacity for flexibly regulating fear and anxiety, the functional architecture of the EAc network, and the degree to which the BST and the Ce are characterized by distinct patterns of functional connectivity, remains incompletely understood.

Building on prior work (**Table 1**), we used a novel combination of approaches to trace and compare the intrinsic functional connectivity of the BST and the Ce. Whole-brain ‘resting-state’ functional MRI (fMRI) data were acquired from a relatively large (*n*=130) sample of psychiatrically healthy, racially diverse, community-dwelling adults, providing increased statistical power and generalizability. Given the challenges of imaging the EAc (A. S. Fox, Oler, Tromp, et al., 2015; Shackman & Fox, 2016), several techniques were used to maximize effective spatial resolution, including a multiband imaging sequence with 2-mm^3^ nominal resolution, boundary-based co-registration (Greve & Fischl, 2009), a novel brain-extraction (‘skull-stripping’) approach, and diffeomorphic normalization (Avants, Epstein, Grossman, & Gee, 2008; Avants et al., 2011; Avants et al., 2010; Klein et al., 2009). To further enhance anatomical specificity, analyses were conducted using spatially unsmoothed data and newly developed, high-precision seeds. Collectively, these techniques enabled us to compare the intrinsic functional connectivity of the BST and the Ce with an unparalleled combination of statistical sensitivity and anatomical precision (**Table 1**). Understanding these functional networks is important: it would provide a baseline against which to compare a range of special populations—including individuals at risk for developing mental illness and patients suffering from psychiatric disorders—and it would inform our understanding of the EAc’s role in normal and pathological fear and anxiety.

**Table 1.**
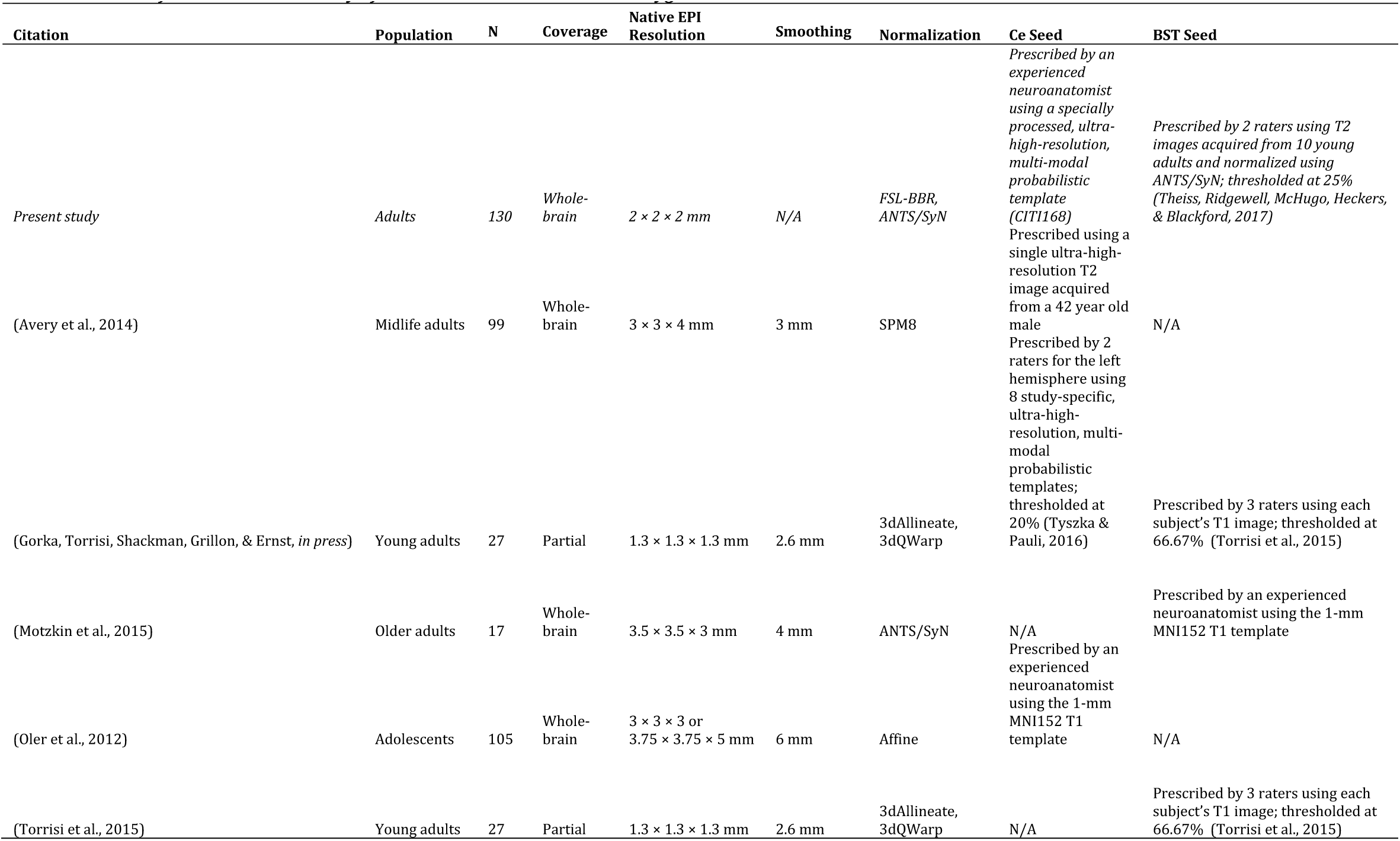
Intrinsic functional connectivity of the human central extended amygdala.

## MATERIALS AND METHODS

### Subjects

Data were extracted from the publicly available Nathan Kline Institute-Rockland Sample (NKI-RS) (<u>http://fcon_1000.projects.nitrc.org/indi/enhanced</u>; Nooner et al., 2012) for 185 adults (18-40 years old). Exclusionary criteria included: positive drug urine screens (*n*=12), current psychiatric diagnosis at the time of the imaging session (*n*=14), incomplete MRI data (*n*=15), and incomplete demographic data (*n*=5). Using procedures detailed below, 18 additional subjects were excluded due to excessive motion artifact (*n*=8), susceptibility artifact (*n*=9), or unusable T1 scans (*n*=1). The final sample consisted of 130 racially (54.6% white, 29.2% African-American, 21.8% other, 10.0% Asian) and ethnically (13.9% Hispanic) diverse subjects (59 males, *M*=25.3 years, *SD*=6.1). A post hoc power analysis performed using G*Power (version 3.1.9.2; Faul, Erdfelder, Buchner, & Lang, 2009; Faul, Erdfelder, Lang, & Buchner, 2007) indicated 98.9% power to detect a ‘medium-sized’ effect (Cohen’s *d*=.50) at two-tailed α=.001 (uncorrected).

### Data Acquisition

MRI data were acquired using a Siemens Magnetom Trio Tim 3 Tesla scanner and 32-channel head-coil (<u>http://fcon_1000.projects.nitrc.org/indi/enhanced/mri_protocol.html</u>). T1-weighted anatomical images were acquired using a magnetization-prepared, rapid-acquisition, gradient-echo sequence (inversion time: 900 ms; repetition time: 1,900 ms; echo time: 2.52 ms; flip angle: 9°; field-of-view: 250 x 250; matrix: 256 x 256; number of slices: 176 sagittal; slice thickness: 1 mm). Building on prior work with partial-brain coverage (Gorka, Torrisi, Shackman, Grillon, & Ernst, *in press;* Torrisi et al., 2015), functional scans were obtained using a T_2_*-weighted echo-planar image (EPI) sequence (multiband acceleration: 4; repetition time: 1,400 ms; echo time: 30 ms; flip angle: 65°; number of excitations: 1; field-of-view: 224 x 224 mm; number of slices: 64 oblique-axial; matrix: 112 x 112; slice thickness: 2mm; gap: ∼0mm; volumes: 404), enabling us to survey the entire brain.

### Data Processing Pipeline

#### Brain extraction and normalization

Given our focus on the BST and the Ce, methods were optimized to minimize spatial normalization error and incidental spatial blurring. Consistent with other work (Acosta-Cabronero, Williams, Pereira, Pengas, & Nestor, 2008; Fein et al., 2006; Fischmeister et al., 2013), unpublished observations by our group demonstrate that the quality of spatial normalization is enhanced by using a brain-extracted (i.e., ‘skull-stripped’ or ‘de-skulled’) template and brain-extracted T1 images. This advantage is particularly evident for publicly available datasets, such as the NKI-RS, where portions of the skull and tissue in the region of the face have been manually removed (‘de-faced’) by the curators to mitigate risks to subject confidentiality (i.e., ‘anonymized’ or ‘de-identified’). However, this benefit is only realized when the quality of the extraction is sufficiently high and consistent, as with images that have been manually extracted by an experienced neuroanatomist. To ensure consistently high-quality extractions, we implemented a multi-tool strategy (for a similar approach, see Najafi, Kinnison, & Pessoa, 2017). For each inhomogeneity-corrected (using N4; Tustison et al., 2014) T1 image, six extraction masks were generated. Five masks were generated using BET (Smith, 2002), BSE (Shattuck, Sandor-Leahy, Schaper, Rottenberg, & Leahy, 2001), 3dSkullstrip (Cox, 1996), ROBEX (Iglesias, Liu, Thompson, & Tu, 2011), and SPM unified segmentation (Ashburner & Friston, 2005), respectively. The sixth mask was generated by applying the inverse spatial transformation (see below) to the MNI152 brain mask distributed with FSL. Specifically, for each subject, the de-faced T1 image was spatially normalized to the MNI152 template using the unified segmentation approach implemented in SPM12; (2) the 1-mm MNI152 template was de-faced to match the idiosyncratic de-facing of the T1 image; (3) the original T1 image was normalized to the individually de-faced 1-mm template using SyN; and (4) the inverse transformation was used to ‘reverse-normalize’ the MNI152 brain mask distributed with FSL to native space. Next, a best-estimate extraction mask was determined by consensus, requiring agreement across four or more extraction techniques. Using this mask, each T1 image was extracted and spatially normalized to the 1-mm MNI152 template using the high-precision diffeomorphic approach implemented in SyN (mutual information cost function; Avants et al., 2008; Avants et al., 2011; Avants et al., 2010; Klein et al., 2009). The average of the 130 normalized T1 images is depicted in **Supplementary Figure S1**.

#### EPI data

The first 3 volumes of each EPI scan were removed and the remaining volumes were de-spiked and slice-time corrected using default settings in AFNI (Cox, 1996). Recent methodological work indicates that de-spiking is more effective than ‘scrubbing’ (Jo et al., 2013; Power, Schlaggar, & Petersen, 2015; Siegel et al., 2014) for attenuating motion-related artifacts in intrinsic functional connectivity. Spike- and slice-time-corrected EPI data were co-registered to the corresponding brain-extracted, native-space T1 image using the boundary-based registration technique implemented in FSL (Greve & Fischl, 2009) and converted to a compatible file format using Convert3d (<u>https://sourceforge.net/p/c3d</u>). Motion correction was then performed using ANTS (<u>https://stnava.github.io/ANTs</u>). The maximum value of the frame-to-frame displacement was calculated for each subject and *z*-transformed. Subjects with a z-score greater than 1.96 (*p*=.05) were excluded (*n*=8). To minimize incidental spatial blurring, the transformation matrices for motion correction, co-registration, and spatial normalization were concatenated and applied to the EPI data in a single step. Normalized EPI data were resampled to 2-mm^3^ voxels using 5^th^–order splines. To maximize spatial resolution, no additional spatial filters were applied, consistent with recent recommendations (Stelzer, Lohmann, Mueller, Buschmann, & Turner, 2014; Turner & Geyer, 2014). Each EPI and T1 dataset was visually inspected before and after processing for quality assurance. To quantify susceptibility artifact in the medial temporal lobe (MTL), we computed the ratio of mean signal in the amygdala relative to the caudate and putamen separately for each hemisphere and subject and then standardized across subjects (i.e., *z*-transformed). Preliminary visual inspection indicated that values greater than ∼2.50 were associated with substantial signal loss (‘drop-out’) in the MTL. Accordingly, subjects with *z*-scores < −2.50 were excluded (*n*=9) (for a similar approach, see Birn et al., 2014). To attenuate physiological noise, white matter (WM) and cerebrospinal fluid (CSF) time-series were identified by thresholding the tissue prior images distributed with FSL, as in prior work by our group (Birn et al., 2014) and others (e.g., Coulombe, Erpelding, Kucyi, & Davis, 2016). The EPI time-series was orthogonalized with respect to the first 3 right eigenvectors of the data covariance matrix from the WM and CSF compartments (Behzadi, Restom, Liau, & Liu, 2007), a Legendre polynomial series (1^st^-5^th^-order), and motion estimates (6 parameters lagged by 0, 1, and 2 volumes), consistent with recent recommendations (Hallquist, Hwang, & Luna, 2013). Orthogonalized time-series were bandpass filtered (0.009-0.10 Hz) using AFNI. Using 3dFWHMx, the mean spatial smoothness of the orthogonalized data was estimated to be ∼2.28 mm^3^.

#### Seed regions

The BST seed was implemented using a previously published probabilistic region of interest thresholded at 25% (Theiss, Ridgewell, McHugo, Heckers, & Blackford, 2017). Building on prior work by our group (Birn et al., 2014; Nacewicz, Alexander, Kalin, & Davidson, 2014; Najafi et al., 2017; Oler et al., 2012; Oler et al., 2017), the Ce was manually prescribed by an experienced neuroanatomist (B.M.N.) using a specially processed version of the CITI168 high-resolution (0.7-mm), multimodal (T1/T2) probabilistic template (http://evendim.caltech.edu/amygdala-atlas; Tyszka & Pauli, 2016) and guided by the atlas of Mai and colleagues (Mai et al., 2007). The methods used for processing the template and prescribing the Ce seed are detailed in the Supplement (**Supplementary Figures S2-S5**). Consistent with prior reports (Birn et al., 2014; Entis, Doerga, Barrett, & Dickerson, 2012; Hrybouski et al., 2016), visual inspection indicated that this approach provides enhanced anatomical sensitivity and selectivity compared to the more widely used centromedial amygdala region-of-interest distributed with FSL (Amunts et al., 2005) (**Supplementary Figure S5**). The BST and Ce seeds are depicted in **Figure 1b** and **Supplementary Figure S6**. To minimize partial volume artifacts, seeds were decimated to the 2-mm MNI template using an iterative procedure that maintained a consistent seed volume across templates. Specifically, each seed was minimally smoothed using a Gaussian kernel and the voxel size was dilated by 0.1-mm and resliced (linear interpolation), enabling us to identify a threshold that approximated the original seed volume and better preserved anatomical boundaries.

### Analytic Plan

We adopted a standard *a priori* seed-based approach to quantifying intrinsic functional connectivity (Biswal, Yetkin, Haughton, & Hyde, 1995; M. D. Fox et al., 2005). For each subject, SPM12 (http://www.fil.ion.ucl.ac.uk/spm/software/spm12) and in-house Matlab code was used to perform a voxelwise regression between the artifact-attenuated, average seed time series and voxel times series throughout the brain. Single-subject regression analyses was performed using the Cochrane-Orcutt procedure for estimating autoregressive error, which is more efficient and potentially less biased than ordinary least-squares (Stocker, 2007). In order to identify regions showing consistent functional connectivity with the BST or Ce seeds across subjects, we tested the intercept in regression models, equivalent to a single-sample *t* test (*p* < .05, whole-brain Šidák corrected; ≥ 80 mm^3^) (Birn et al., 2014; Oler et al., 2010; Šidák, 1967). A minimum conjunction (Boolean ‘AND’) was used to identify regions showing significant coupling with both seeds (Nichols, Brett, Andersson, Wager, & Poline, 2005) and a paired *t*-test was used to assess differential functional connectivity. For ease of interpretation, differential connectivity was only examined in the subset of 12,004 voxels where functional connectivity was significant for one or both seeds (*p* < .05, Šidák corrected for the 12,004 voxel region-of-interest; ≥ 80 mm^3^). This approach circumvents the need to interpret significant differences (e.g., BST > Ce) in regions where neither seed shows significant functional connectivity. As an additional check on the integrity of the data and our approach, we confirmed our ability to identify the default mode network (**Supplementary Figure S7**). Figures were created using MRIcron (http://people.cas.sc.edu/rorden/mricron).

## RESULTS

### Subcortical Regions

As shown in **Figure 2** and **Supplementary Figure S8**, whole-brain regression analyses revealed robust coupling between the BST and the Ce regions (*p*<.05, whole-brain Šidák corrected; **Tables 2-4**). Analyses seeded in the BST showed significant functional connectivity with neighboring regions of the basal forebrain and basal ganglia as well as distal voxels in the region of the Ce. The complementary pattern was observed for the Ce seed—significant functional connectivity with neighboring regions of the dorsal amygdala and with distal voxels located in the region of the BST. Consistent with invasive tracing studies (Oler et al., 2017), the BST and Ce also showed robust coupling with anatomically intermediate voxels located in the SLEA, the ribbon of subcortical gray matter (‘substantia innominata’) encompassing the ventral amygdalofugal pathway (**Figure 3**). Finally, both seeds showed significant functional connectivity with the anterior hippocampus (**Figure 2**).

**Figure 2.**
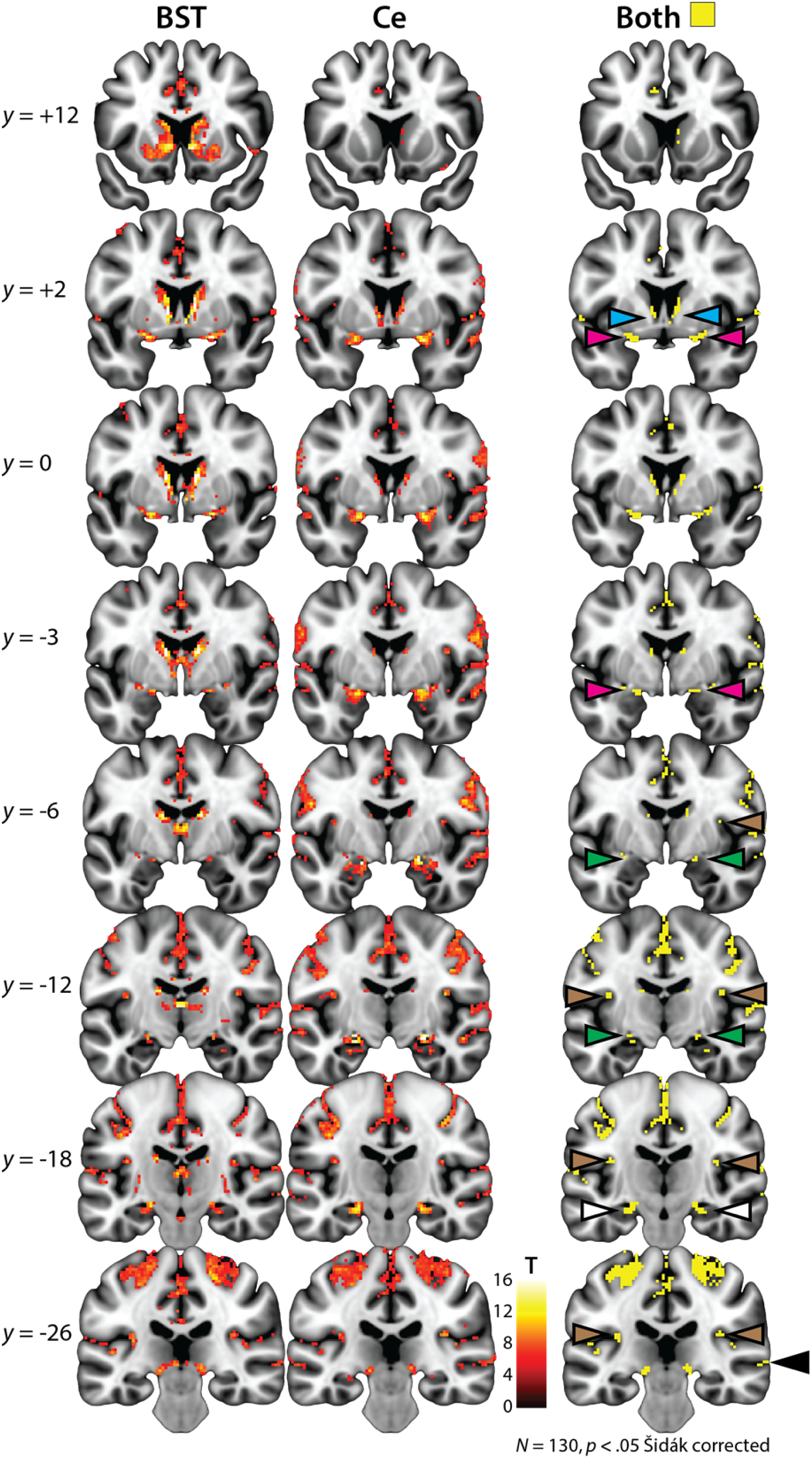
Intrinsic functional connectivity of the EAc. Left and center columns depict the results of whole-brain regression analyses for the BST and the Ce seed regions, respectively, conservatively thresholded at *p*<.05 whole-brain Šidák corrected. The right column depicts the intersection or conjunction (Boolean ‘AND’) of the two thresholded maps (Nichols et al., 2005). The BST seed showed significant functional connectivity with neighboring voxels in the basal forebrain (*cyan* arrowheads) as well as voxels in the region of the Ce (*green* arrowheads), while the Ce seed showed significant coupling with neighboring voxels in the dorsal amygdala as well as distal voxels in the region of the BST. Analyses also demonstrated that the BST and Ce exhibit robust functional connectivity with intermediate voxels located along the path of the ventral amygdalofugal pathway in the sublenticular extended amygdala (*magenta* arrowheads). Finally, both regions showed significant coupling with the anterior hippocampus (*white* arrowheads), posterior insula (*brown* arrowheads), and superior temporal sulcus (*black* arrowheads). Note: Results are depicted here and reported in the accompanying tables for clusters of at least 80 mm^3^. Abbreviations—BST, bed nucleus of the stria terminalis; Ce, central nucleus of the amygdala; EAc, central division of the extended amygdala; L, left hemisphere; R, right hemisphere.

**Figure 3.**
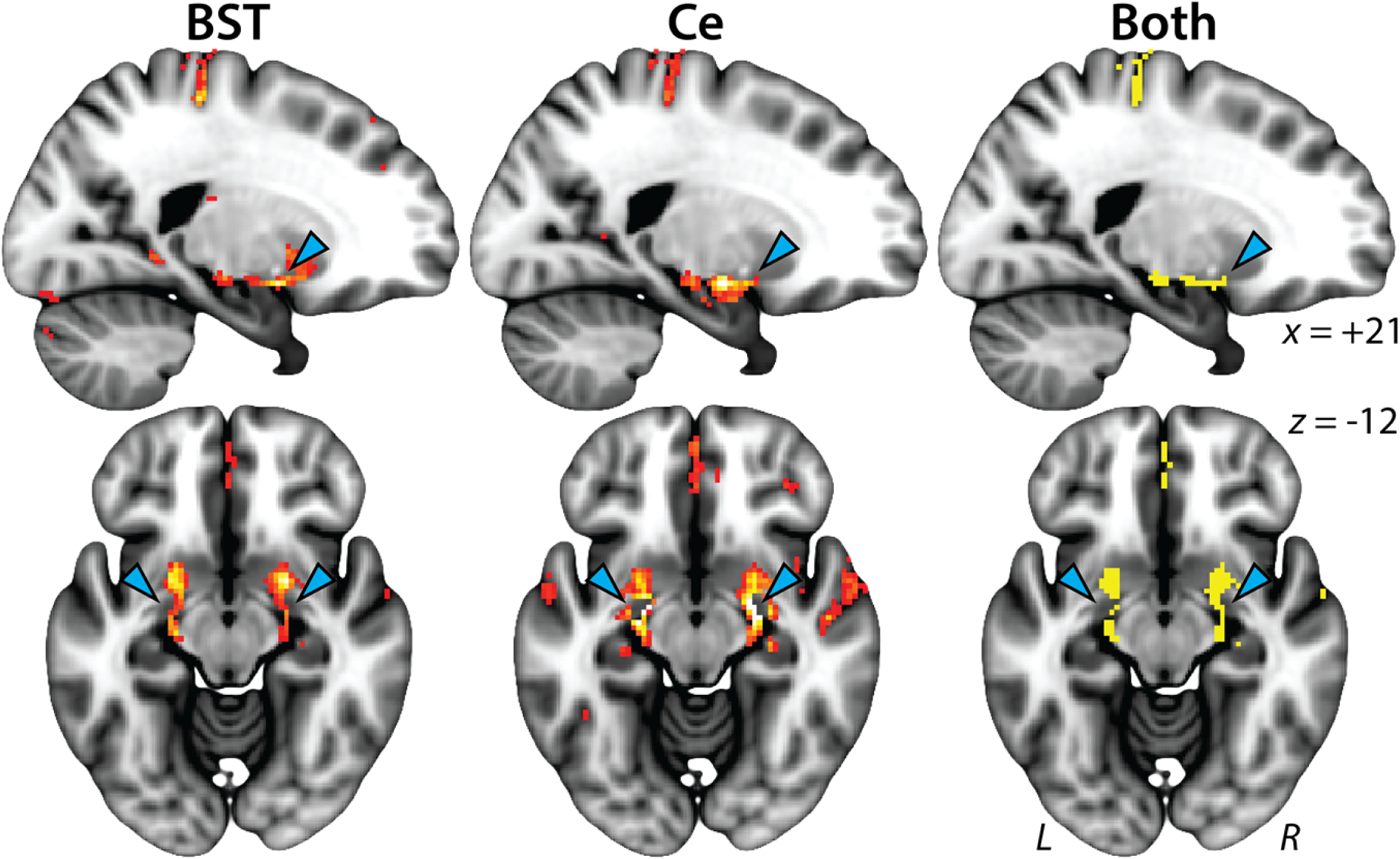
The BST and the Ce are functionally linked via the SLEA. Clusters in the region of the SLEA (*cyan* arrowheads). Conventions are similar to **Figure 2**. Abbreviations—BST, bed nucleus of the stria terminalis; Ce, central nucleus of the amygdala; L, left hemisphere; R, right hemisphere; SLEA, sublenticular extended amygdala.

**Table 2.**
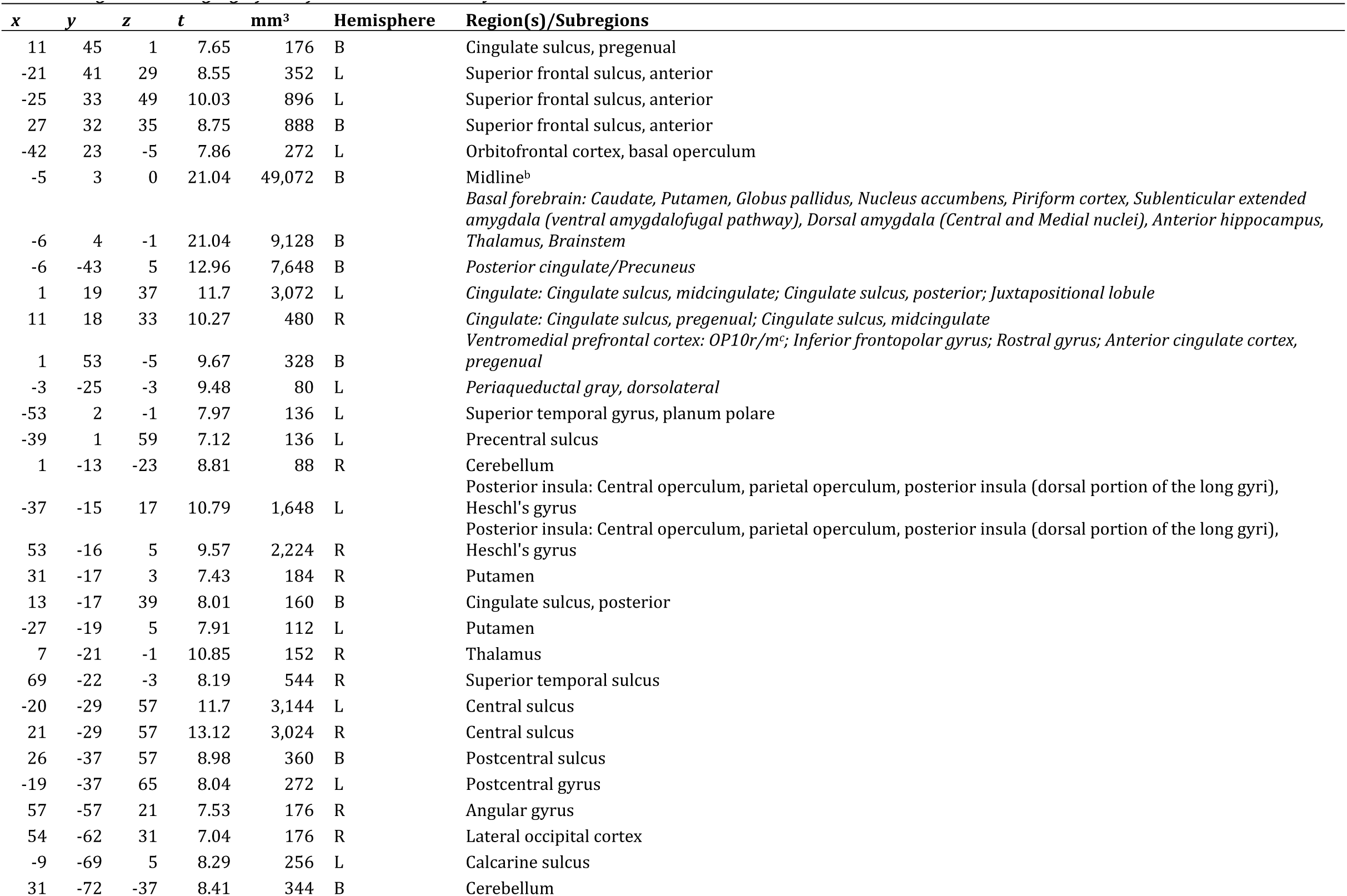
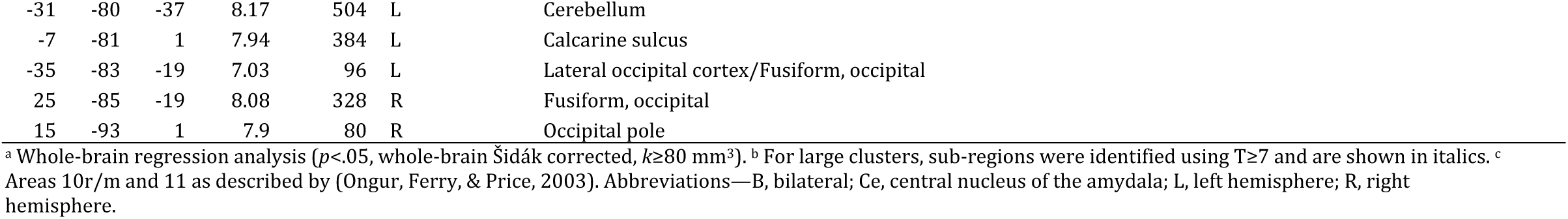
Regions showing significant functional connectivity with the BST^a^.

**Table 3.**
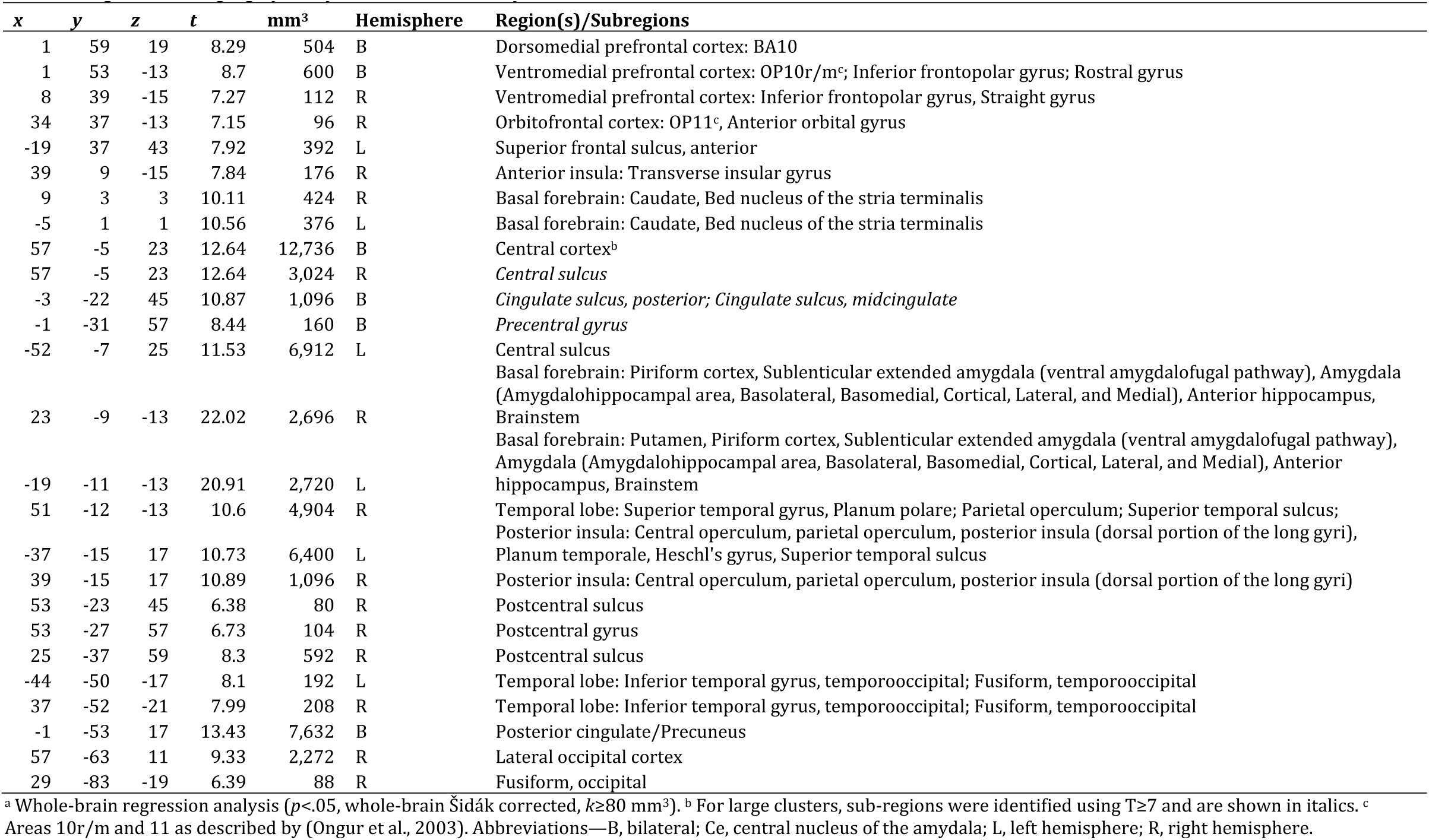
Regions showing significant functional connectivity with the Ce^a^.

**Table 4.** Regions showing significant functional connectivity with both the BST and the Ce^a^.

| **X** | **y** | **z** | **mm<sup>3</sup>** | **Hemisphere** | **Region(s)/Subregions** |
| --- | --- | --- | --- | --- | --- |
| 1 | 61 | 21 | 48 | R | Dorsomedial prefrontal cortex: BA10 |
| 3 | 59 | 17 | 16 | R | Dorsomedial prefrontal cortex: BA10 |
| 1 | 57 | 13 | 24 | R | Dorsomedial prefrontal cortex: BA10 |
| 1 | 53 | 19 | 80 | R | Dorsomedial prefrontal cortex: BA10 |
| -1 | 49 | 27 | 24 | L | Dorsomedial prefrontal cortex: BA10 |
| 1 | 39 | -15 | 296 | R | Ventromedial prefrontal cortex: OP10r/m <sup>b</sup>; Inferior frontopolar gyrus; Rostral gyrus |
| -21 | 27 | 37 | 304 | L | Superior frontal sulcus, anterior |
| 55 | 7 | -3 | 8 | R | Temporal pole |
| 63 | 7 | -1 | 664 | R | Planum temporale |
| 9 | 5 | -1 | 384 | R | Basal forebrain: Caudate, Bed nucleus of the stria terminalis |
| -9 | 5 | 35 | 3,448 | L | Cingulate: Cingulate sulcus, posterior midcingulate; Cingulate sulcus, posterior |
| -5 | 3 | -1 | 312 | L | Basal forebrain: Caudate, Bed nucleus of the stria terminalis |
| 5 | 1 | -3 | 8 | R | Bed nucleus of the stria terminalis |
| -53 | 1 | -1 | 40 | L | Planum polare |
| 53 | 1 | -1 | 24 | R | Planum polare |
| -1 | 1 | 47 | 8 | L | Juxtapositional lobule |
| -17 | -3 | -15 | 376 | L | Dorsal amygdala: Amygdalohippocampal area, Central, Cortical, Medial |
| 63 | -3 | 17 | 2,640 | R | Central sulcus |
| 61 | -5 | -13 | 392 | R | Superior temporal sulcus |
| 29 | -11 | -23 | 976 | R | Hippocampus |
| -41 | -15 | 31 | 2,648 | L | Central sulcus |
| 5 | -15 | 73 | 8 | R | Precentral gyrus |
| 63 | -17 | -7 | 8 | R | Superior temporal sulcus |
| -53 | -17 | 9 | 8 | L | Heschl's gyrus |
| -21 | -19 | -17 | 616 | L | Hippocampus/Dorsal amygdala: Basolateral, Basomedial, Central, Medial |
| -57 | -19 | 9 | 152 | L | Planum temporale |
| 13 | -19 | 39 | 40 | R | Cingulate sulcus, posterior |
| 3 | -19 | 67 | 16 | R | Precentral gyrus |
| -47 | -25 | 3 | 880 | L | Planum temporale |
| 47 | -25 | 7 | 728 | R | Planum temporale |
| -25 | -31 | 67 | 96 | L | Postcentral gyrus |
| 3 | -33 | 49 | 16 | R | Posterior cingulate |
| 27 | -37 | 55 | 232 | R | Postcentral sulcus |
| -21 | -39 | 63 | 128 | L | Postcentral gyrus |
| 3 | -39 | 63 | 8 | R | Postcentral gyrus |
| 11 | -53 | 1 | 6,792 | R | Posterior cingulate/Precuneus |
| 55 | -57 | 19 | 136 | R | Angular gyrus |
| 45 | -59 | 29 | 8 | R | Lateral occipital cortex |
| 31 | -85 | -19 | 88 | R | Occipital fusiform |

<sup>a</sup> Minimum conjunction (Boolean 'AND') analysis ( $p < .05$ , whole-brain Šidák corrected, $k \geq 80$ mm<sup>3</sup>). <sup>b</sup> Area 10r/m as described by (Ongur et al., 2003). Abbreviations—B, bilateral; BST, bed nucleus of the stria terminalis; Ce, central nucleus of the amygdala; L, left hemisphere; R, right hemisphere.

Compared to the Ce, the BST showed significantly stronger coupling with several subcortical regions, including the basal ganglia (i.e., nucleus accumbens, caudate, and putamen), thalamus, and the brainstem in the region of the dorsal periaqueductal gray (PAG) (**Figure 4**, **Supplementary Figure S9**, and **Table 5**). The only subcortical regions showing stronger functional connectivity with the Ce were located in the anterior hippocampus and dorsal amygdala, and included the amygdalohippocampal area and basolateral, basomedial, cortical, and medial nuclei.

**Figure 4.**
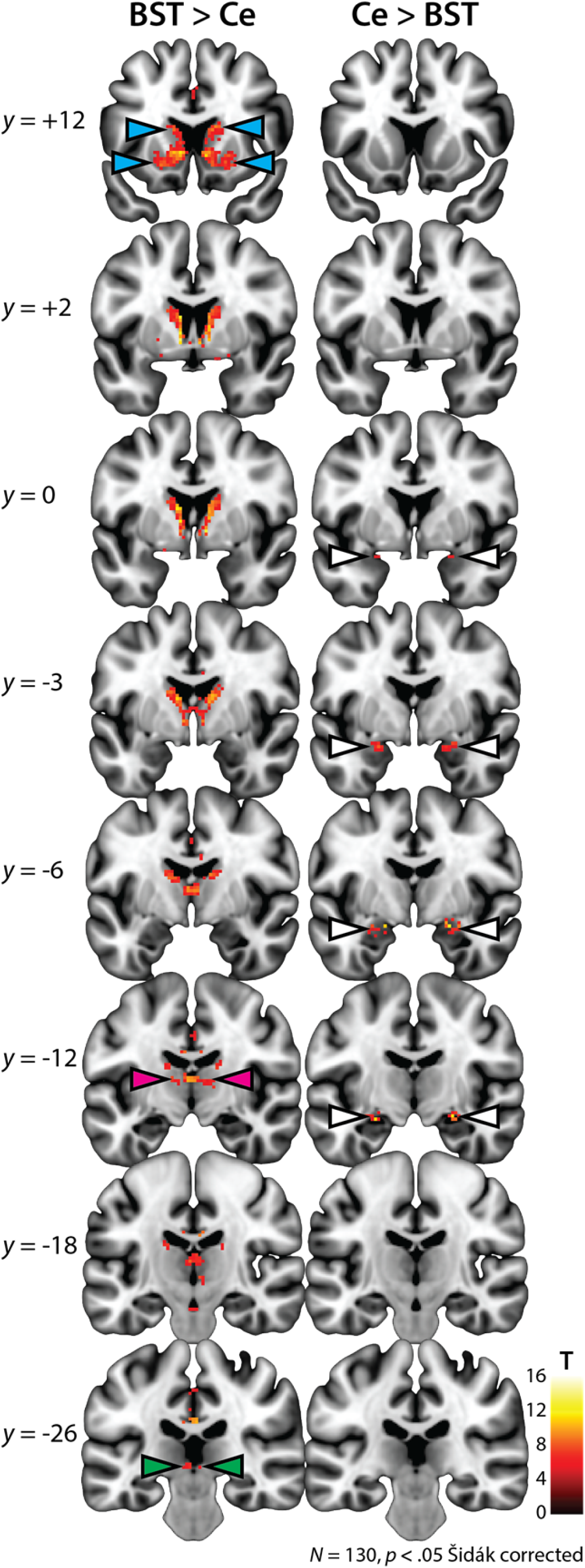
Differential functional connectivity of the BST vs. Ce. Results of a paired *t*-test comparing the intrinsic functional connectivity of the BST and Ce. The left and right columns depict regions showing significantly stronger coupling with the BST and Ce, respectively. For ease of interpretation, differences were only examined in the subset of 12,004 voxels (2-mm^3^) where functional connectivity was significant for the BST, the Ce, or both seeds (see **Figures 2-3**). Consistent with other analyses, results were thresholded at *p*<.05 Šidák corrected for the extent of the 12,004-voxel mask. Results revealed significantly stronger coupling between the BST and the basal ganglia, including the caudate, putamen, and nucleus accumbens (*cyan* arrowheads). The BST also showed significantly stronger connectivity with the thalamus (*magenta* arrowheads) and a region of the brainstem consistent with the dorsal periaqueductal gray (*green* arrowheads; see also **Supplementary Figure S9**). The only regions showing stronger connectivity with the Ce were neighboring regions of the amygdala (*white* arrowheads), including voxels in the region of the amygdalohippocampal area and the basolateral, basomedial, cortical, and medial nuclei. Note: Results are depicted here and reported in the accompanying tables for clusters of at least 80 mm^3^. Abbreviations—BST, bed nucleus of the stria terminalis; Ce, central nucleus of the amygdala; L, left hemisphere; R, right hemisphere.

**Table 5.**
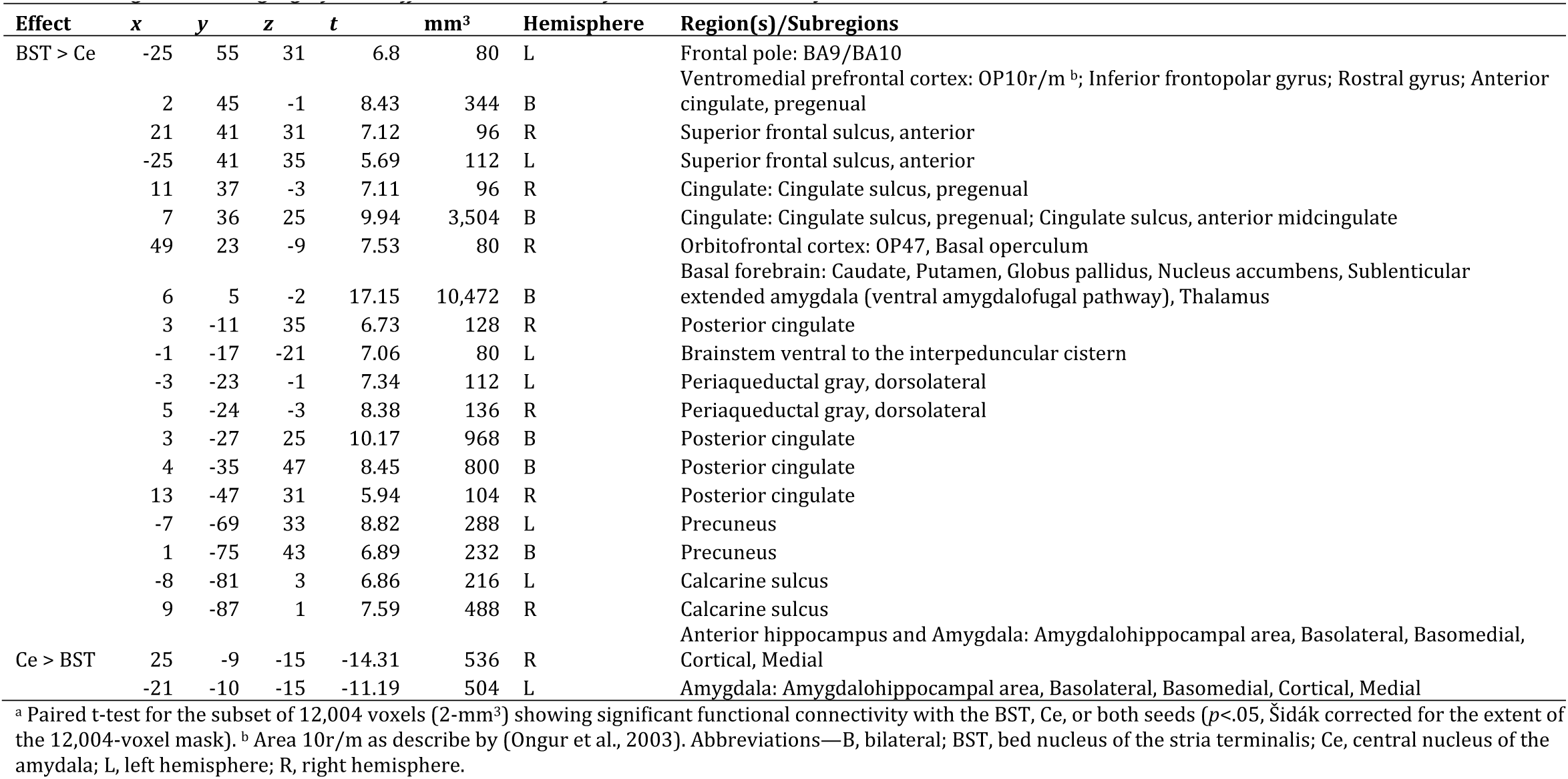
*Regions showing significant differences in intrinsic functional connectivity between the BST and the Ce^a^*.

### Cortical Regions

As shown in **Figures 2** and **5**, the BST and the Ce showed significant functional connectivity with several cortical regions, including the ventromedial prefrontal cortex (vmPFC), posterior MCC, posterior insula, posterior cingulate/precuneus, and parts of the ventral visual processing stream (e.g., superior temporal sulcus, fusiform cortex) (**Tables 2-4**). As shown in **Figure 5**, relative to the Ce, the BST displayed significantly stronger coupling with a cluster centered on the anterior MCC that extends into the pregenual anterior cingulate cortex (pgACC) and vmPFC (**Figure 5, far-right panels, and Table 5**). As detailed in the Supplement (**Supplementary Figure S10**), control analyses indicated that these effects could not be attributed to regional differences in signal quality, as indexed by several widely used metrics (e.g., the temporal signal-to-noise ratio [tSNR]).

**Figure 5.**
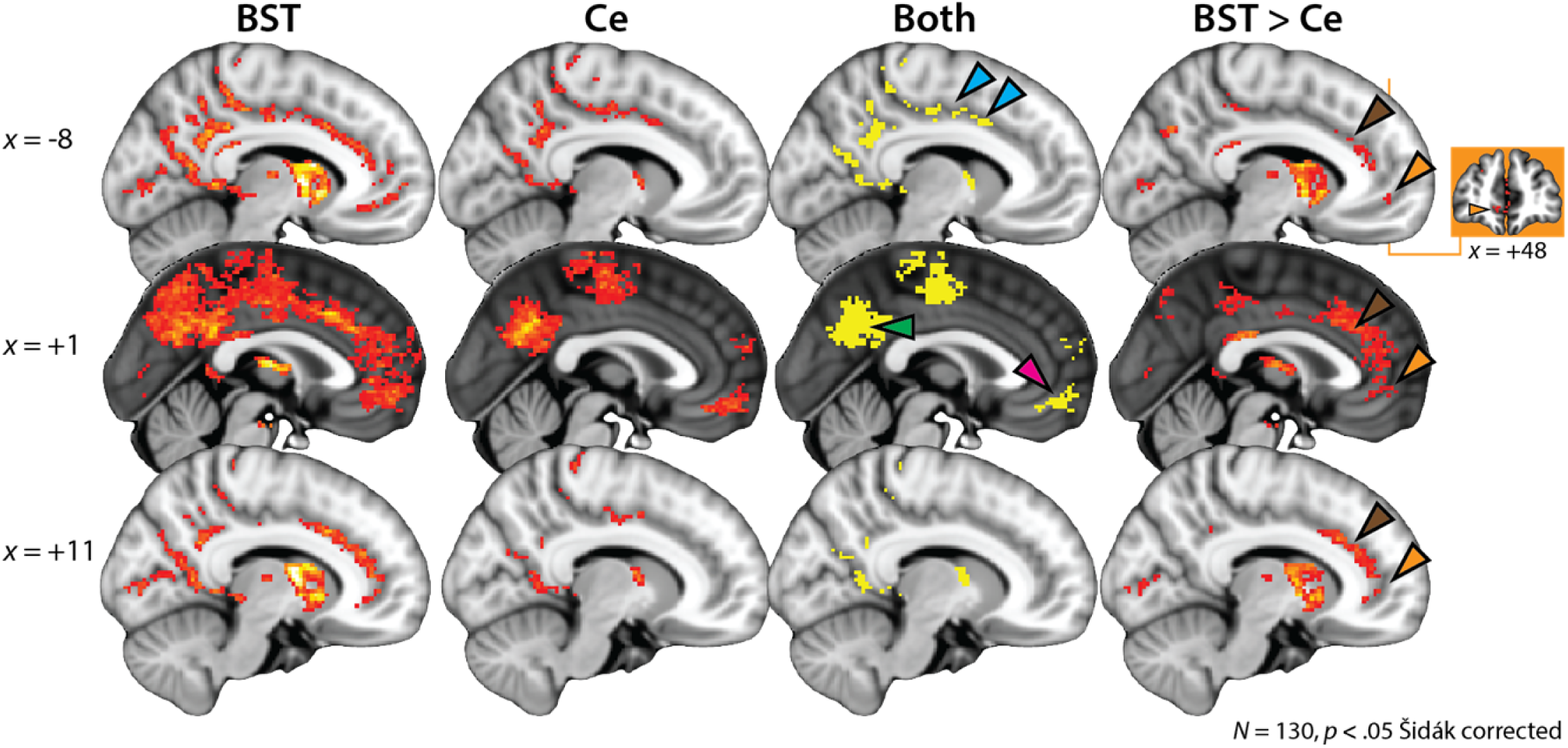
Intrinsic functional connectivity of the EAc and midline cortical regions. The first two columns depict the results of whole-brain regression analyses for the BST and Ce seed regions, respectively (*p*<.05, whole-brain Šidák corrected). The third column depicts the intersection (Boolean ‘AND’) of the two thresholded maps. The fourth column depicts the results of a paired *t*-test comparing the intrinsic functional connectivity of the BST and Ce (*p*<.05, small-volume Šidák corrected). Both seeds show significant functional connectivity with the posterior cingulate/precuneus (*green* arrowhead), posterior MCC (*cyan* arrowheads), and vmPFC (*magenta* arrowhead). Relative to the Ce, the BST shows significantly stronger coupling with the anterior MCC and pgACC (*brown* arrowheads) as well as the vmPFC (*orange* arrowheads). Orange inset depicts a coronal slice through the vmPFC cluster, which extends along the rostral-caudal axis from area 10r/m and the inferior frontopolar gyrus to the rostral gyrus and pgACC. Conventions are similar to **Figure 2** (first three columns) and **Figure 4** (fourth column). Abbreviations—BST, bed nucleus of the stria terminalis; Ce, central nucleus of the amygdala; EAc, central divisions of the extended amygdala; L, left hemisphere; MCC, midcingulate cortex; pgACC, pregenual anterior cingulate cortex; R, right hemisphere; vmPFC, ventromedial prefrontal cortex.

## DISCUSSION

The EAc plays a central role in assembling states of fear and anxiety and is implicated in the development, maintenance, and recurrence of a range of debilitating psychiatric disorders. The present findings provide new insights into the normative architecture of the EAc functional network. Our results indicate that the BST and the Ce are robustly interconnected via the SLEA (**Figure 3** and **Supplementary Figure S8**), consistent with anatomical and functional tracing studies in monkeys (Birn et al., 2014; Oler et al., 2012; Oler et al., 2017). By and large, the BST and the Ce showed patterns of functional connectivity that were similar to one another and concordant with prior human imaging research (**Table 6**). Both regions showed significant coupling with subcortical and cortical regions implicated in fear and anxiety—including the anterior hippocampus, insula, MCC, and vmPFC (**Figures 2 and 5**)—reinforcing the hypothesis that these regions represent a functionally coherent macro-circuit (Alheid & Heimer, 1988; A. S. Fox, Oler, Tromp, et al., 2015; Oler et al., 2012; Shackman & Fox, 2016).

**Table 6.** Intrinsic functional connectivity of the EAc across human imaging studies^b,f^.

| Seed | Citation | NAcc | Cd | Putamen | GP | BST | SLEA | Amygdala | Hippocampus | Thalamus | PAG | vmPFC/<br>OFC | pgACC | MCC | Insula | Precuneus |
| --- | --- | --- | --- | --- | --- | --- | --- | --- | --- | --- | --- | --- | --- | --- | --- | --- |
| BST | *Present study* | + | + | + | + | N/A | + | + | + | + | + | + | + | + | +<sup>p</sup> | + |
|  | (Avery et al., 2014)<sup>c</sup> | + | + | + | + | N/A |  | + | + | + |  | + | + | + | +<sup>a,p</sup> | + |
|  | (Torrissi et al., 2015) | + | + | + |  | N/A | + | + | + | + | + | + | + |  | +<sup>p</sup> | + |
|  |  | 3/3 | 3/3 | 3/3 | 2/3 | N/A | 2/3 | 3/3 | 3/3 | 3/3 | 2/3 | 3/3 | 3/3 | 2/3 | 3/3 | 3/3 |
| Ce | *Present study* |  | + |  |  | + | + | + | + |  |  | + |  | + | +<sup>a,p</sup> | + |
|  | (Gorka et al., *in press*)<sup>d</sup> | + | + | + |  | + | + | + | + | + | + | + | + |  | +<sup>a,p</sup> | + |
|  | (Oler et al., 2012)<sup>e</sup> | + |  | + |  | + |  | + | + | + |  |  |  |  | + |  |
|  |  | 2/3 | 2/3 | 2/3 | 0/3 | 3/3 | 2/3 | 3/3 | 3/3 | 2/3 | 1/3 | 2/3 | 1/3 | 2/3 | 3/3 | 2/3 |

Despite their many similarities, it is unlikely that the BST and the Ce are completely interchangeable (A. S. Fox & Shackman, *under review;* Shackman & Fox, 2016). Indeed, the BST showed significantly stronger connectivity with anterior cortical regions (anterior MCC, pgACC and vmPFC), with the posterior cingulate/precuneus, with the medial temporal lobe (striatum and SLEA), and with the brainstem in the region of the dorsal PAG (**Supplementary Figure S9**), whereas the Ce showed stronger connectivity with neighboring regions of the amygdala and anterior hippocampus (**Figures 4-5**)—observations that largely align with recent high-resolution fMRI research (Gorka et al., *in press*) (cf. **Table 1**). Supplementary analyses indicated that these effects were not a consequence of regional differences in signal quality (e.g., tSNR).

We also observed significant coupling between the BST, the Ce, and the vmPFC (i.e., inferior frontopolar gyrus, rostral gyrus, and area OP10), although this effect was stronger for the BST seed region (**Figure 5**). This pattern is consistent with other work (Gorka et al., *in press; their Figure 2e*) and is particularly interesting in light of several recent observations in nonhuman primate models of fear and anxiety. First, intrinsic coupling between the Ce and vmPFC co-varies with the intensity of defensive behaviors and neuroendocrine activity elicited by exposure to human intruder threat in monkeys (Birn et al., 2014). Second, metabolic activity in the Ce, BST, and vmPFC, as well as the anterior hippocampus and PAG, co-varies with these same anxiety-related responses (A. S. Fox, Oler, Shackman, et al., 2015). Third, vmPFC lesions have been shown to reduce these defensive responses and imaging research suggests that this anxiolytic effect is likely to be mediated by ‘downstream’ alterations in BST metabolism (A. S. Fox et al., 2010; Kalin, Shelton, & Davidson, 2007; Motzkin et al., 2015; Rudebeck, Saunders, Prescott, Chau, & Murray, 2013). These and other observations (e.g., Grayson et al., 2016; Kalin et al., 2016; Kalin et al., 2004) motivate the hypothesis that fear and anxiety partially reflect a core neural system encompassing the BST, Ce, vmPFC, anterior hippocampus, and PAG (A. S. Fox, Oler, Shackman, et al., 2015; Oler et al., 2016; Shackman, Tromp, et al., 2016).

Our results revealed evidence of robust coupling between the BST, Ce, and rostral cingulate and they hint at a rostro-caudal gradient: both seeds showed coupling with the posterior MCC, while the BST showed significantly stronger coupling with a cluster centered on the anterior MCC (**Figure 5**). Notably, the MCC and a region consistent with the BST are frequently co-activated in imaging studies of Pavlovian fear conditioning (Fullana et al., 2016; Mechias, Etkin, & Kalisch, 2010) and uncertain threat anticipation (Alvarez et al., 2011; Alvarez et al., 2015; J. M. Choi et al., 2012; Grupe et al., 2013; Herrmann et al., 2016; Klumpers et al., 2015; McMenamin et al., 2014; Somerville et al., 2010). We have previously hypothesized that the MCC uses information about pain, negative feedback, punishment, and threat to bias responding in situations where the optimal course of action is uncertain or risky (Cavanagh & Shackman, 2015; Shackman et al., 2011) (see also de la Vega et al., 2016) and the present results highlight the potential importance of communication between the MCC and the EAc, particularly the BST, for this kind of top-down control. A key challenge for future research will be to more formally characterize the nature of task-related interactions among these three key regions using graph-theoretic or related analytic techniques (McMenamin et al., 2014; Najafi et al., 2017).

Clearly, a number of other important challenges remain. As with most brain imaging studies, our analyses do not permit mechanistic inferences and like other studies focused on functional connectivity, our conclusions are tempered by questions about the origins and significance of correlated fluctuations in the blood-oxygen-level dependent (BOLD) fMRI signal (Akam & Kullmann, 2014; Cabral, Kringelbach, & Deco, 2014; Logothetis, 2008). A key challenge for future research will be to use a combination of mechanistic (e.g., optogenetic) and whole-brain imaging techniques to clarify the specific causal contributions of the regions highlighted here and more precisely delineate the nature of their functional interactions (A. S. Fox & Shackman, *under review;* Shackman & Fox, 2016; Wiegert, Mahn, Prigge, Printz, & Yizhar, 2017).

Existing treatments for anxiety disorders are inconsistently effective or associated with significant adverse effects (Bystritsky, 2006; Cloos & Ferreira, 2009), highlighting the need to identify and understand the neural mechanisms controlling the experience and expression of fear and anxiety. Building on prior mechanistic and imaging research, the present study indicates that the BST and the Ce are marked by broadly similar patterns of intrinsic functional connectivity, with both regions showing significant coupling with the EAc, anterior hippocampus, insula, MCC, and vmPFC. Despite these similarities, the BST displayed significantly stronger connectivity with the rostral cingulate and vmPFC. These observations provide a baseline against which to compare a range of special populations—including individuals at risk for developing mental illness and patients suffering from psychiatric disorders—and inform our understanding of the role of the EAc in normal and pathological fear and anxiety. The use of a relatively large sample increases our confidence in the robustness of these results (Button et al., 2013; A. S. Fox, Lapate, Davidson, & Shackman, *in press;* Poldrack et al., 2017). Finally, from a methodological perspective, these results highlight the value of several new techniques for EAc seed prescription and image registration/normalization. The former is likely to be useful for other investigators focused on the BST and Ce, while the latter will be advantageous for any investigator confronted with the problem of spatially normalizing structural images that have been modified—anatomically ‘anonymized’ or ‘de-identified’—prior to public release (e.g., Holmes et al., 2015; Nooner et al., 2012).

## CONTRIBUTIONS

R.M.T, A.J.S., and J.F.S. designed the study. M.D.S. coordinated data extraction. B.M.N. developed and implemented the protocol for segmenting the Ce seed and the HyperEdge method. J.F.S. developed and implemented the novel image registration/normalization pipeline. R.M.T. and J.F.S. processed data. J.F.S. and A.J.S. analyzed data. R.M.T., A.J.S., and A.S.F. interpreted data. R.M.T., A.J.S., A.S.F., and B.M.N. wrote the paper. A.J.S., R.M.T., B.M.N., and A.S.F. created figures. R.M.T. and A.J.S. created tables. S.T. provided theoretical guidance. A.J.S. funded and supervised all aspects of the study. All authors contributed to reviewing and revising the paper and approved the final version.

## ACKNOWLEDGEMENTS

Authors acknowledge assistance from J. Blackford, K. DeYoung, L. Friedman, M. Milham, and D. Tromp and critical feedback from N. Fox, L. Pessoa, and E. Redcay. We also wish to thank Drs. R. Poldrack and A. Holmes for guidance on the signal quality analyses. This work was supported by the University of California, Davis; University of Maryland, College Park; University of Wisconsin—Madison; and National Institutes of Health (DA040717 and MH107444). Authors declare no conflicts of interest.

## DATA AVAILABILITY/SHARING

Upon acceptance of this manuscript, all of the key statistical maps will be uploaded to NeuroVault.org. Raw data are publicly available (<u>http://fcon_1000.projects.nitrc.org/indi/enhanced/</u>).

## Supplementary method and results to accompany—

### Spatial Normalization

Given our focus on the BST and the Ce, methods were optimized to minimize spatial normalization error and incidental spatial blurring. Unpublished observations by our group demonstrate that the quality of spatial normalization is enhanced by using a brain-extracted (i.e., ‘skull-stripped’ or ‘de-skulled’) template and brain-extracted T1 images, consistent with prior reports (Acosta-Cabronero, Williams, Pereira, Pengas, & Nestor, 2008; Fein et al., 2006; Fischmeister et al., 2013). This advantage is particularly evident for publicly available datasets, such as the NKI-RS, where portions of the skull and tissue in the region of the face have been manually removed (‘de-faced’) by the curators to mitigate risks to subject confidentiality. However, this benefit is only realized when the quality of the extraction is sufficiently high and consistent, as with images that have been manually extracted by a well-trained neuroanatomist. To ensure consistently high-quality extractions, we implemented a multi-tool strategy (for a similar approach, see Najafi, Kinnison, & Pessoa, 2017). For each inhomogeneity-corrected (using N4; Tustison et al., 2014) T1 image, six extraction masks were generated. Five masks were generated using BET (Smith, 2002), BSE (Shattuck, Sandor-Leahy, Schaper, Rottenberg, & Leahy, 2001), 3dSkullstrip (Cox, 1996), ROBEX (Iglesias, Liu, Thompson, & Tu, 2011), and SPM unified segmentation (Ashburner & Friston, 2005), respectively. The sixth mask was generated by applying the inverse spatial transformation (see below) to the MNI152 brain mask distributed with FSL^1^. Next, a best-estimate extraction mask was determined by consensus, requiring agreement across four or more extraction techniques. Using this mask, each T1 image was extracted and spatially normalized to the 1-mm MNI152 template using the diffeomorphic approach implemented in SyN (mutual information cost function; Avants, Epstein, Grossman, & Gee, 2008; Avants et al., 2011; Avants et al., 2010), the most accurate normalization tool (Klein et al., 2009). The average of the resulting normalized T1 images (*n*=130) is depicted in **Supplementary Figure S1**.

**Supplementary Figure S1.**
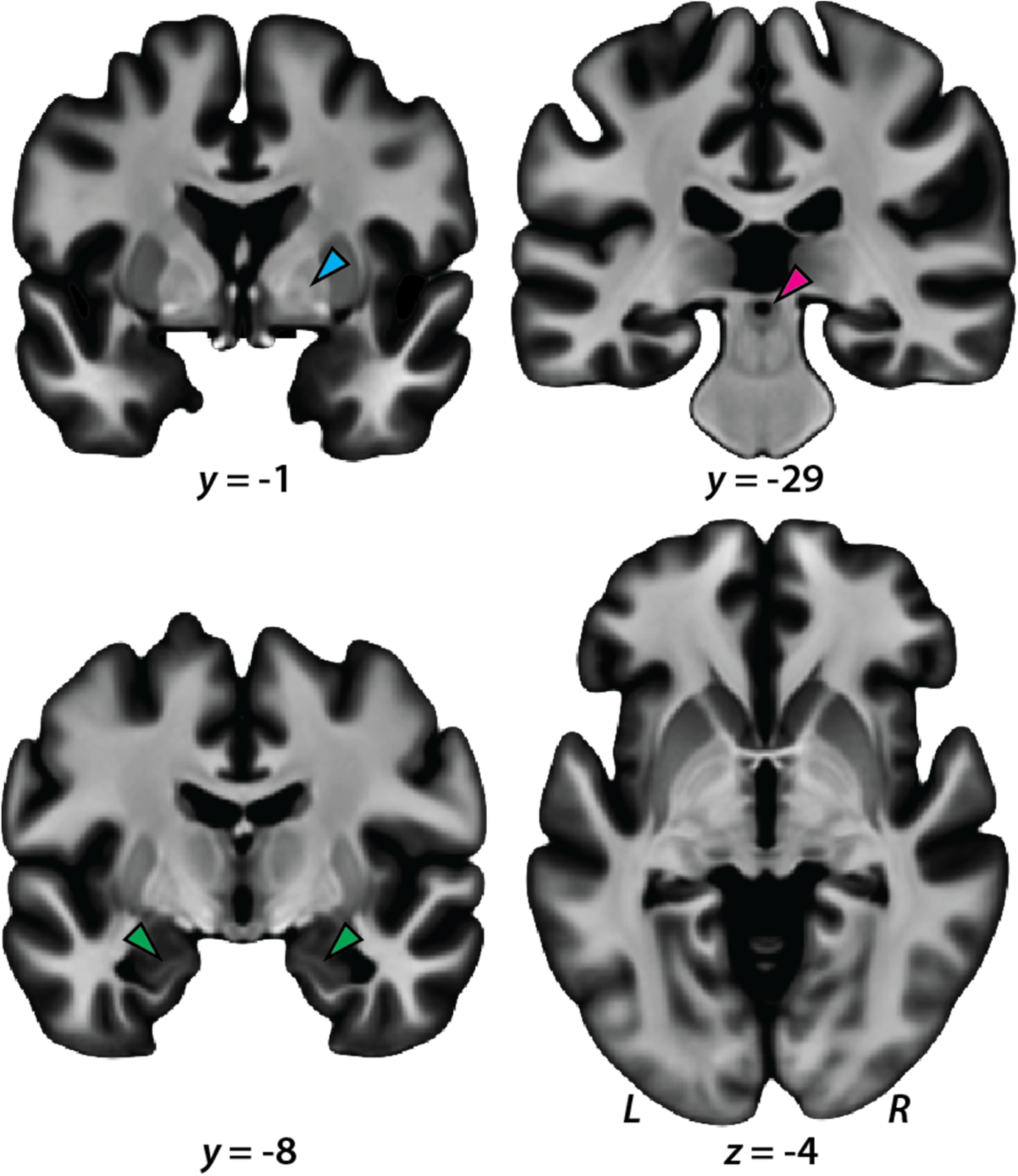
Mean normalized T1 image. Figure depicts representative slices from the average of the 130 diffeomorphically normalized T1 images. Note the preservation of fine detail in the medial medullary lamina of the globus pallidus (*cyan* arrowhead), periaqueductal gray (*magenta* arrowhead), and alveus (*green* arrowheads).

### Ce Seed

#### Overview

Building on prior work by our group using similar methods (Birn et al., 2014; Nacewicz, Alexander, Kalin, & Davidson, 2014; Najafi et al., 2017; Oler et al., 2012; Oler et al., 2017), the Ce was manually prescribed by an experienced neuroanatomist (B.M.N.) based on the atlas of Mai and colleagues (Mai, Paxinos, & Voss, 2007; Prevost, McCabe, Jessup, Bossaerts, & O’Doherty, 2011) using a specially processed version of the CITI168 high-resolution (0.7-mm), multimodal (T1/T2) probabilistic template (http://evendim.caltech.edu/amygdala-atlas; Tyszka & Pauli, 2016). The procedures used for processing the template and prescribing the Ce seed are detailed below.

#### Template processing and co-registration

To maximize acutance (i.e., perceived sharpness) and enable reliable discrimination of Ce boundaries, we implemented a novel edge-detection approach (**Supplementary Figure S2**). Floating point precision was used for all computations. Preliminary work indicated that conventional edge-detection approaches (e.g., Laplacian filtering, AFNI’s *3dedge3* tool) were inadequate. Subsequent testing indicated that the ratio of the 1^st^ and 2^nd^ derivative of spatial intensity differences, which can be conceptualized as a hyperbolically-exaggerated edge map (‘HyperEdge’), provided a sensitive means of detecting anatomical edges in typical T1 and T2 anatomical images. To overcome noise amplification (‘speckle’ artifact)—a key obstacle for edge detection tools—the mean absolute slope across nearest-neighbors in each of the 3 cardinal directions, excluding the intensity of the voxel-of-interest, was computed using histogram-normalized images. Further enhancement was achieved using a variant of the approach described by Srivastava and colleagues (Tucker, Wu, & Srivastava, 2013; Wu & Srivastava, 2014). This enabled us to generate edge maps that could be dynamically thresholded or ‘tune’ to reveal anatomical boundaries that could not otherwise be visually discerned in the template.

More specifically, an edge image X’ (1) is calculated as the square-root of the mean absolute slope across (but not including X_0_) in each of *k* directions. As shown in (2), X’ is then divided by the square-root mean slope of X’ (cf. Cheng, Dryden, & Huang, 2016; Kurtek, 2017) to generate hyperbolic exaggeration of inflection points, that is, a HyperEdge map.

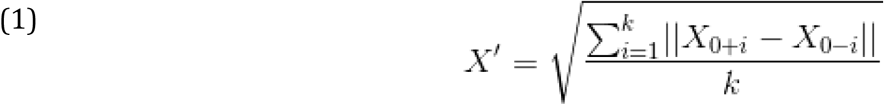

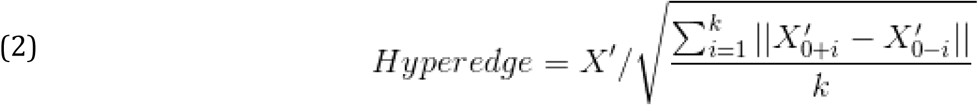

**Supplementary Figure S2.**
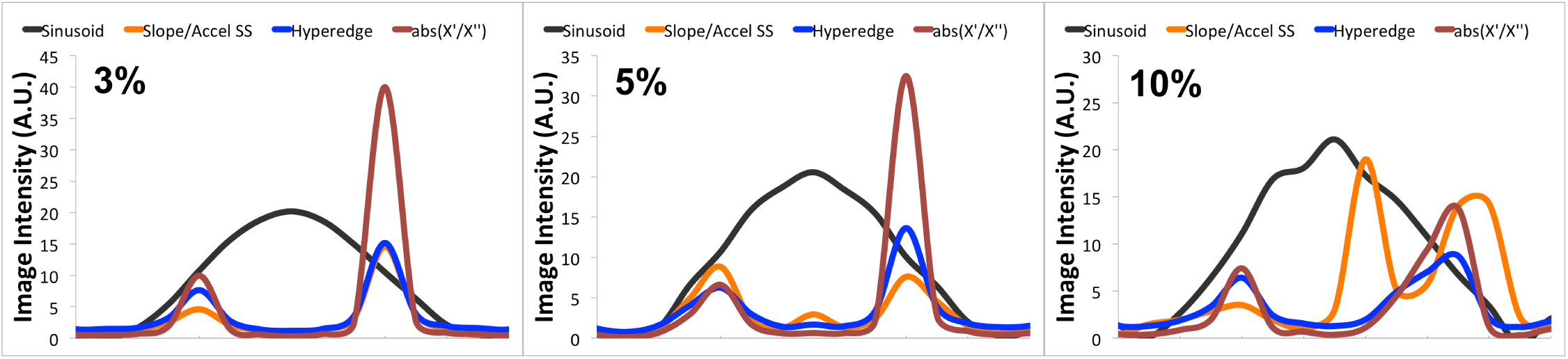
HyperEdge preserves symmetric boundaries even in high noise. In each panel, the profile of an anatomical element (e.g., a thin section of gray matter) is simulated as a sinusoid (*black*) with added white noise of 3%, 5% and 10% for the *left*, *middle* and *right* panels, respectively. A hyperbolic inflection point map using a simple sum of squared differences of the first derivative (Slope; similar to average of absolue-valued Laplacian components) and second derivative (Accel) shows low signal to noise overall (Slope/Accel SS, *orange lines*) and is prone to false peaks at the plateau of a distribution (middle) and at large point deviations (right). Taking the average absolute valued slope across nearest neighbors but excluding the voxel of interest (abs X’/X”, *red lines*) protects against large fluctuations from single-voxel noise (middle and right panels), but inflection point estimates are quickly exaggerated and asymmetric with very slight noise. Taking the square-root of the absolute slope (hyperedge, *blue lines*) greatly reduces effects of noise, consistent with formal analyses in the space of square-root slope in functional data analysis, and produces largely symmetric edge gradients even with very high noise. This robust edge-detection allows coregistration and segmentation of subtle anatomical features with low signal-to-noise.

The Ce seed was prescribed bilaterally using an adapted version of the CITI168 probabilistic template (<u>http://evendim.caltech.edu/amygdala-atlas</u>, version 1.0.1; CIT168_T1w_700um_MNI.nii and CIT168_T2w_700um_MNI.nii). 3dQwarp was used to co-register and up-sample (0.35-mm) the T1 and T2 templates. As shown in **Supplementary Figure S3**, the HyperEdge approach was used to create a dynamically tunable tracing overlay, revealing inter-nuclear boundaries that were not readily apparent in the unprocessed template.

**Supplementary Figure S3.**
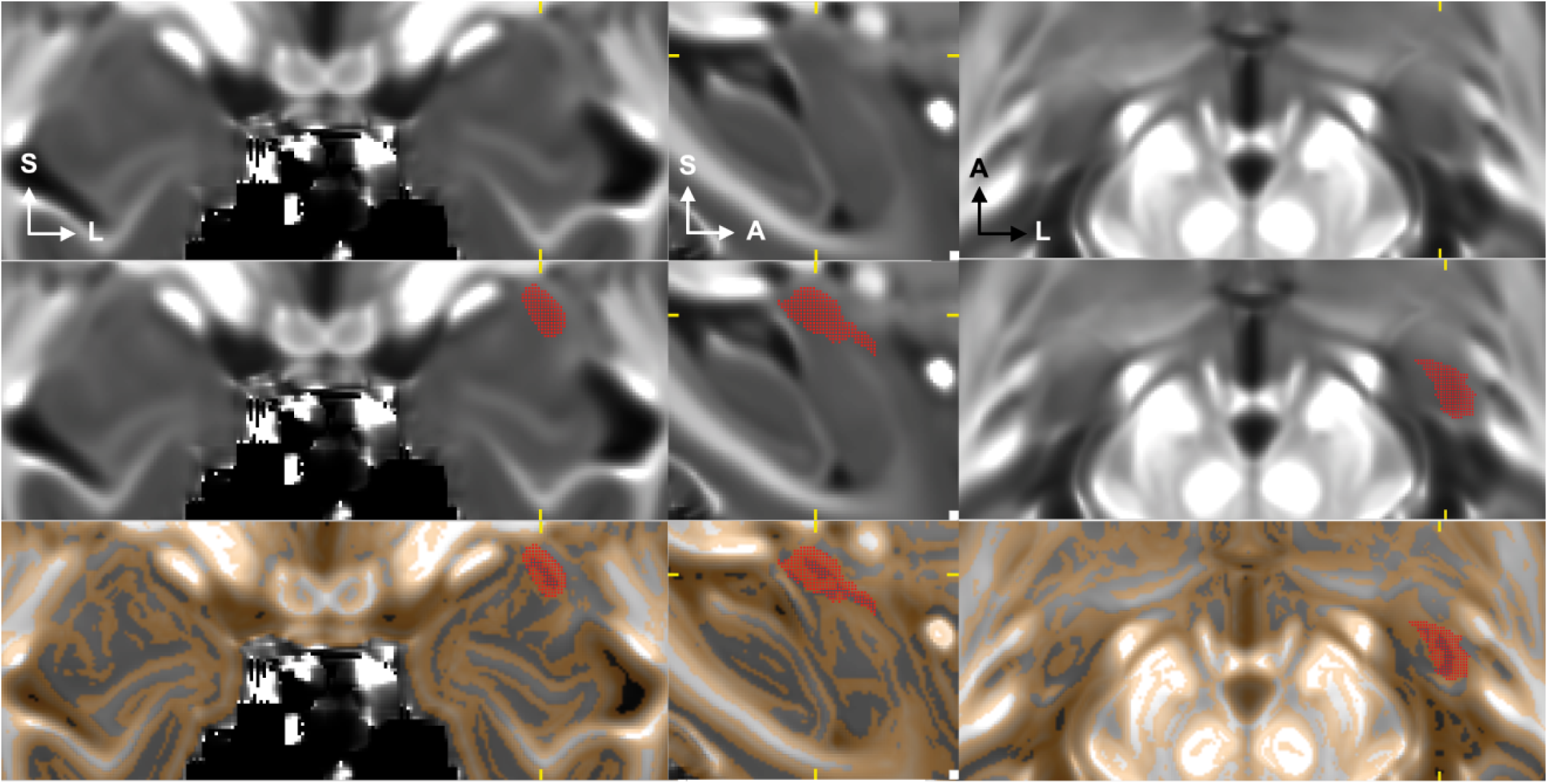
Using the HyperEdge approach to guide Ce prescription. The left Ce seed (*red; middle and bottom rows*) is depicted at a single location in the CITI168 0.35-mm template in the coronal (*left*), sagittal (*middle*), and axial (*right*) planes. The dynamically tunable HyperEdge map is shown in the bottom row (*gold*). Abbreviations—A, anterior; L, left hemisphere; S, superior.

#### Ce protocol

The criteria used for manually prescribing the Ce seed represent an extension of our previously published protocol (Nacewicz et al., 2014; Nacewicz et al., 2006) (for applications, see Chung, Worsley, Nacewicz, Dalton, & Davidson, 2010; Nacewicz et al., 2006) and leverages the additional contrast afforded by the high-resolution, multimodal template and HyperEdge processing technique (**Supplementary Figure S4**). In contrast to other recent work by our group (Birn et al., 2014; Oler et al., 2012) and others (Tyszka & Pauli, 2016), the Ce was prescribed in both the left and right hemispheres. The criteria were derived from the atlas of Mai and colleagues (2007) and hinged on identifying the lateral division of the Ce (CeL) at its first appearance caudally and including surrounding tissue up to the boundary with the ventral putamen (laterally and dorsally) and the more T1-intense basolateral nuclei (ventrally). Moving rostrally, a thin, notch-like band of white matter separates the dorsal portions of the basolateral and lateral nuclei from the Ce. The ventromedial tip of the white matter separating the Ce from the basolateral nuclei was then followed in a straight line to the lateral margin of the optic tract or the rhinal sulcus to form the ventromedial border. A major landmark is the disappearance of the head of the hippocampus, at which point the CeL can no longer be discerned. The Ce curves medially and ventrally during the progression from caudal to rostral slices, and in the sections rostral to the disappearance of the hippocampus, care was taken not to include the peri-amygdalar claustrum (lateral to the Ce). In the middle and rostral slices, portions of the boundary between the Ce and medial nuclei was not evident in the HyperEdge-enhanced T1 and T2 templates. In these cases, the visible portions of the boundary were extrapolated using straight lines. Preliminary traces were refined in all three cardinal planes. In the case of conflicting traces, the axial and coronal slices were favored over the more variable sagittal slice.

**Supplementary Figure S4.**
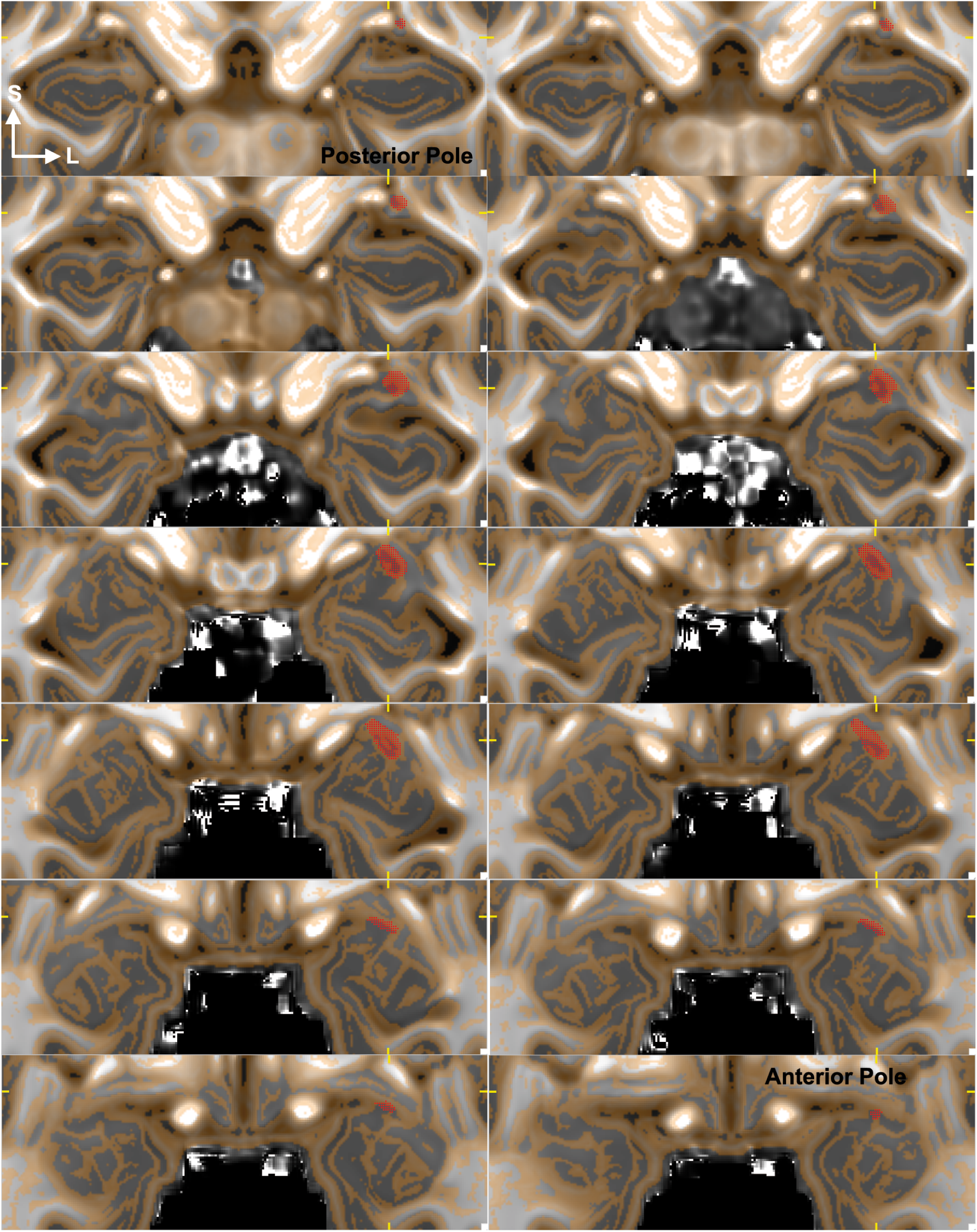
Ce seed in native space. Coronal montage depicts the left Ce seed (*red*) at every third slice. Slices are arranged from posterior (*upper left*) to anterior (*bottom right*). Conventions are described in the legend for **Supplementary Figure S3.**

#### Seed decimation

The resulting high-resolution (0.35-mm) Ce seeds were normalized to an upsampled version of the MNI152 template using 3dQwarp. To minimize partial volume artifacts, left and right Ce seeds were decimated to the 2-mm MNI152 grid using an iterative procedure that maintained a consistent seed volume across templates. Specifically, each seed was minimally smoothed using a Gaussian kernel and the voxel size was dilated by 0.1-mm and resliced (linear interpolation), enabling us to identify a threshold that approximated the original seed volume.

**Supplementary Figure S5.**
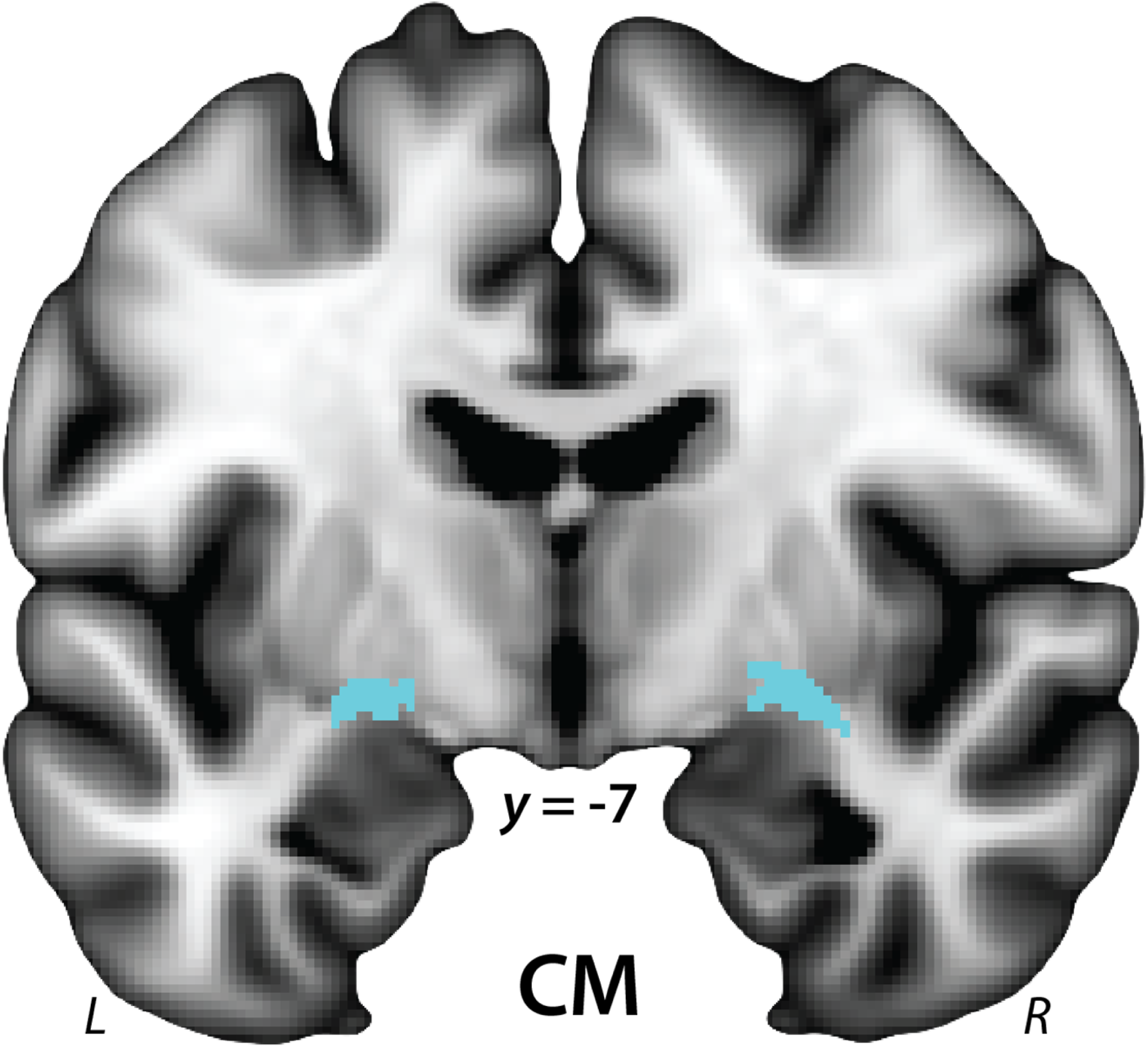
Jülich Centromedial Amygdala (CM) Seeds. The derivation of the widely used probabilistic CM seed (*cyan*) is described in more detail in (Amunts et al., 2005). This figure depicts the version of the CM seed distributed with the FSL software package. The seed has been thresholded at 25% and overlaid on the nonlinear 1-mm MNI152 anatomical template. It is clear that the CM seeds encompass a substantial volume of extra-amygdalar tissue, including regions of white matter, globus pallidus, and putamen. A similar pattern was evident when the seeds were thresholded at 50%. This likely reflects a registration error when the CM seed was normalized to the MNI152 nonlinear template prior to distribution with the FSL software package (Simon Eichoff, *personal communication, 12/15/2016*). For illustrative purposes, 1-mm seeds are shown. Abbreviations—CM, centromedial amygdala; L, left hemisphere; R, right hemisphere.

**Supplementary Figure S6.**
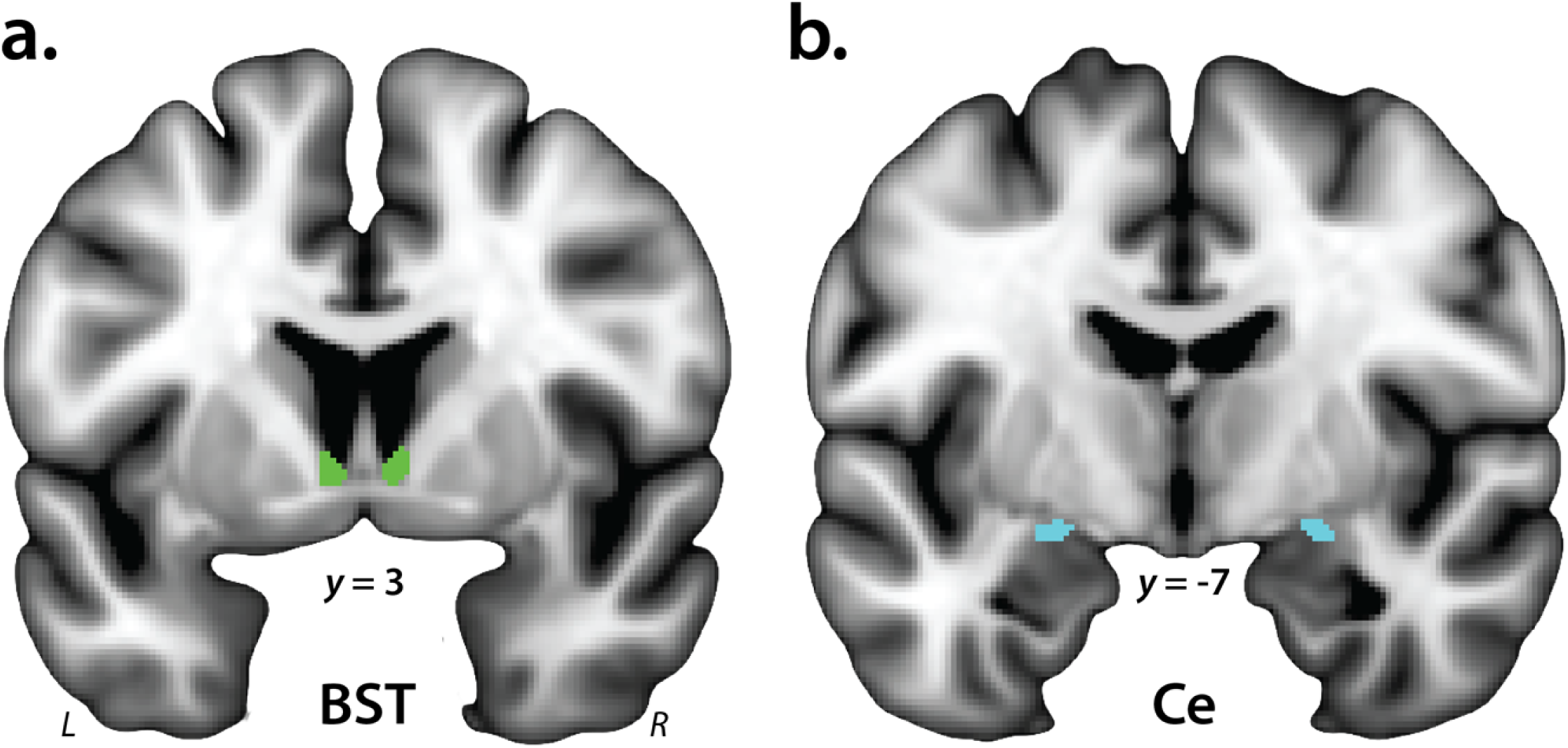
BST and Ce Seeds. ***a. BST seed.*** The derivation of the probabilistic BST seed (*green*) is described in more detail in Theiss and colleagues (2016) and was thresholded at 25%. The seed mostly encompasses the supra-commissural BST, given the difficulty of reliably discriminating the borders of regions below the anterior commissure on the basis of T1-weighted MRI (cf. Kruger, Shiozawa, Kreifelts, Scheffler, & Ethofer, 2015). ***b. Ce seed.*** For illustrative purposes, 1-mm seeds are shown. Analyses employed seeds decimated to the 2-mm resolution of the EPI data. Single-subject l data were visually inspected to ensure that the seeds were correctly aligned to the spatially normalized T1 images. Abbreviations—BST, bed nucleus of the stria terminalis; Ce, central nucleus of the amygdala; L, left hemisphere; R, right hemisphere.

**Supplementary Figure S7.**
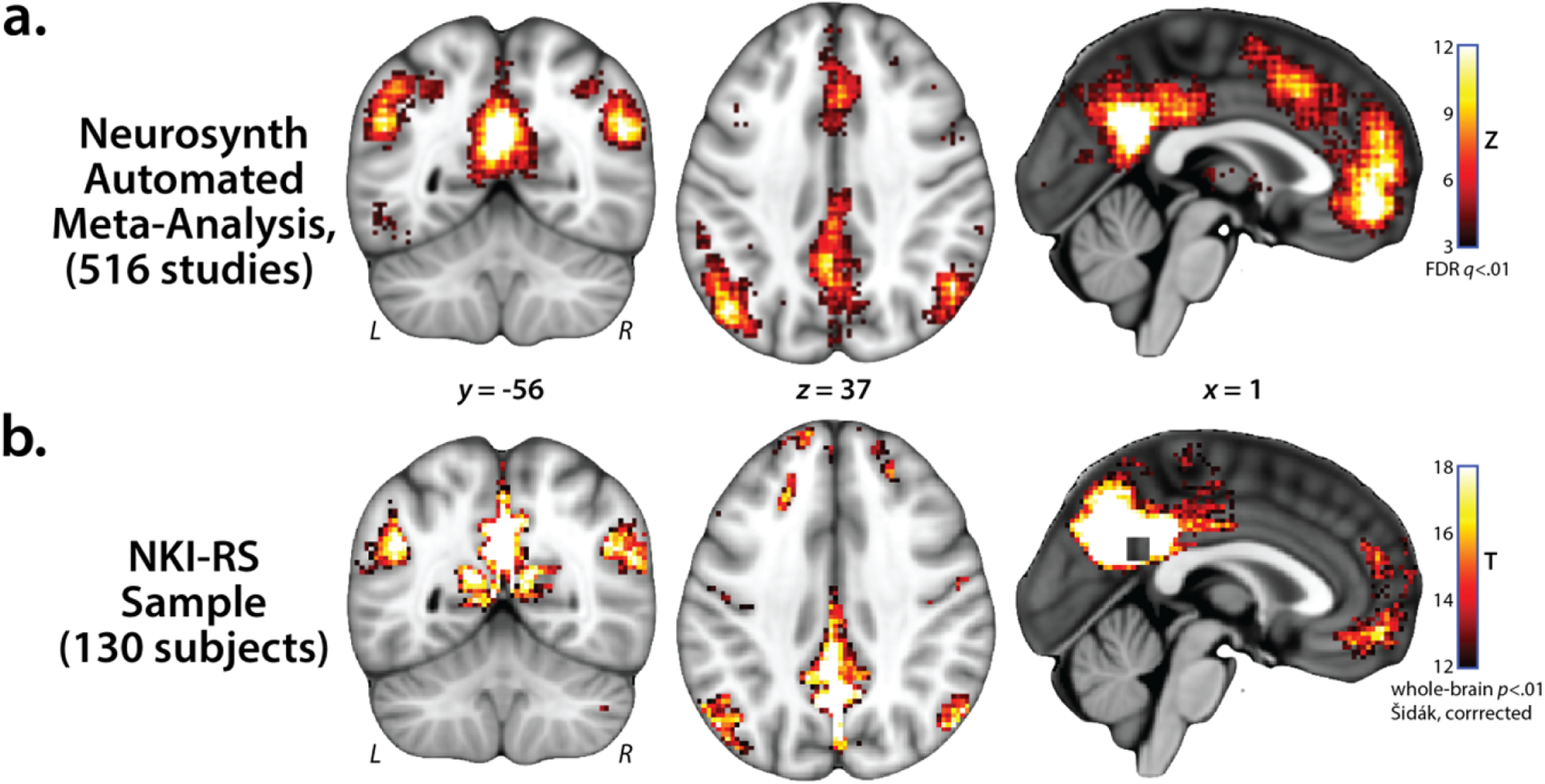
Confirmatory Analysis of the Default Mode Network (DMN). For quality assurance purposes, we performed a confirmatory analysis of the DMN and compared it to an automated meta-analysis of ‘default mode’ performed using Neurosynth (whole-brain FDR *q* < .01) (Yarkoni, Poldrack, Nichols, Van Essen, & Wager, 2011). Our confirmatory analysis was performed using a 10-mm seed (square-shaped region in panel b) centered on the location (*x* = 0, *y* = −50, *z* = 28) in the precuneus showing the strongest reverse-inference association with ‘default mode’ in the Neurosynth database. For illustrative purposes, the resulting map was conservatively thresholded (*t* >16.0, *p* < 9.6 × 10^-23^, uncorrected). As expected, both the automated meta-analysis (panel a) and confirmatory analysis (panel b) revealed regions typical of the DMN, including the posterior cingulate cortex, medial prefrontal cortex, and lateral temporoparietal cortex. Abbreviations—L, left hemisphere; NKI-RS, Nathan Kline Institute-Rockland Sample; R, right hemisphere.

**Supplementary Figure S8.**
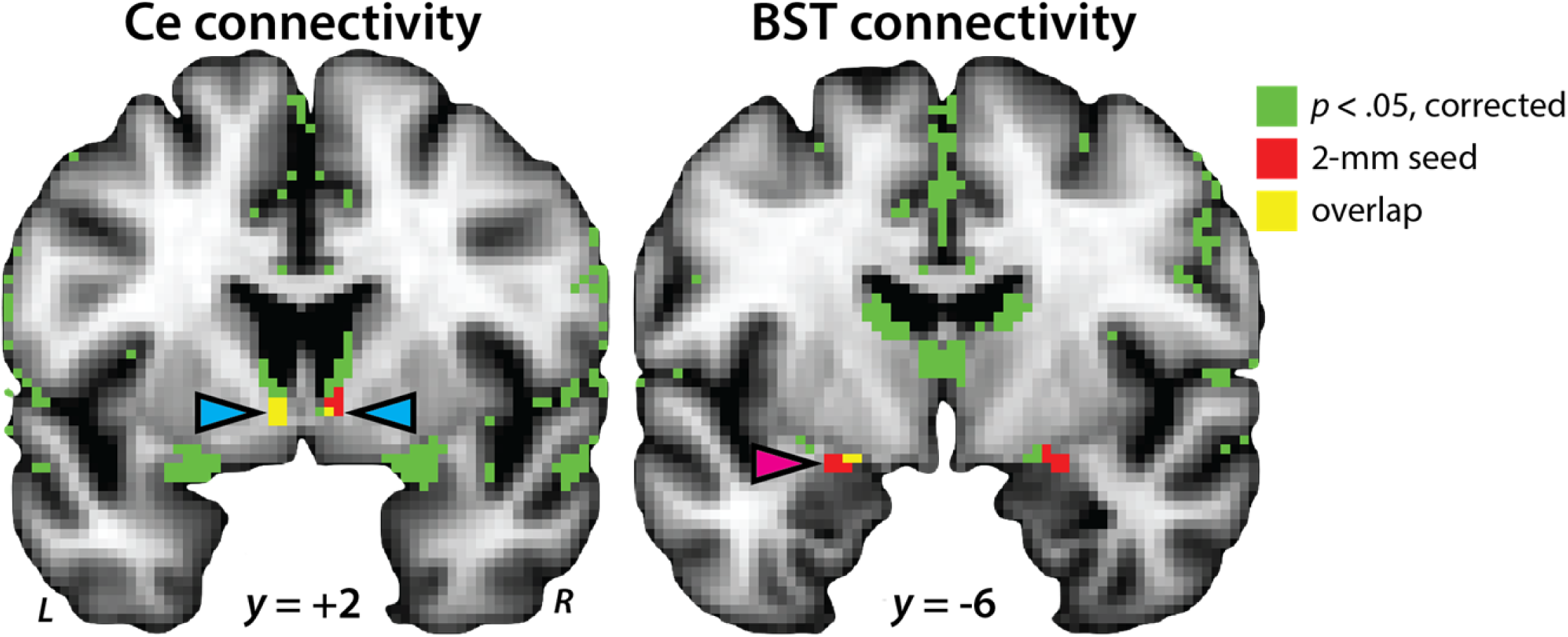
Whole-brain regression analyses revealed robust coupling between the BST and the Ce. Analyses seeded in the Ce showed significant functional connectivity (*p*<.05, whole-brain Šidák corrected; *green*) with voxels located in the region of the BST seed (*cyan* arrowheads; overlap depicted in *yellow*), while analyses seeded in the BST showed significant functional connectivity with voxels located in the region of the Ce seed (*magenta* arrowheads; overlap depicted in *yellow*). For maximal precision, the uninterpolated statistical maps and the seeds are displayed on the 2-mm MNI152 grid used for all analyses. Abbreviations—BST, bed nucleus of the stria terminalis; Ce, central nucleus of the amygdala; L, left hemisphere; R, right hemisphere.

**Supplementary Figure S9.**
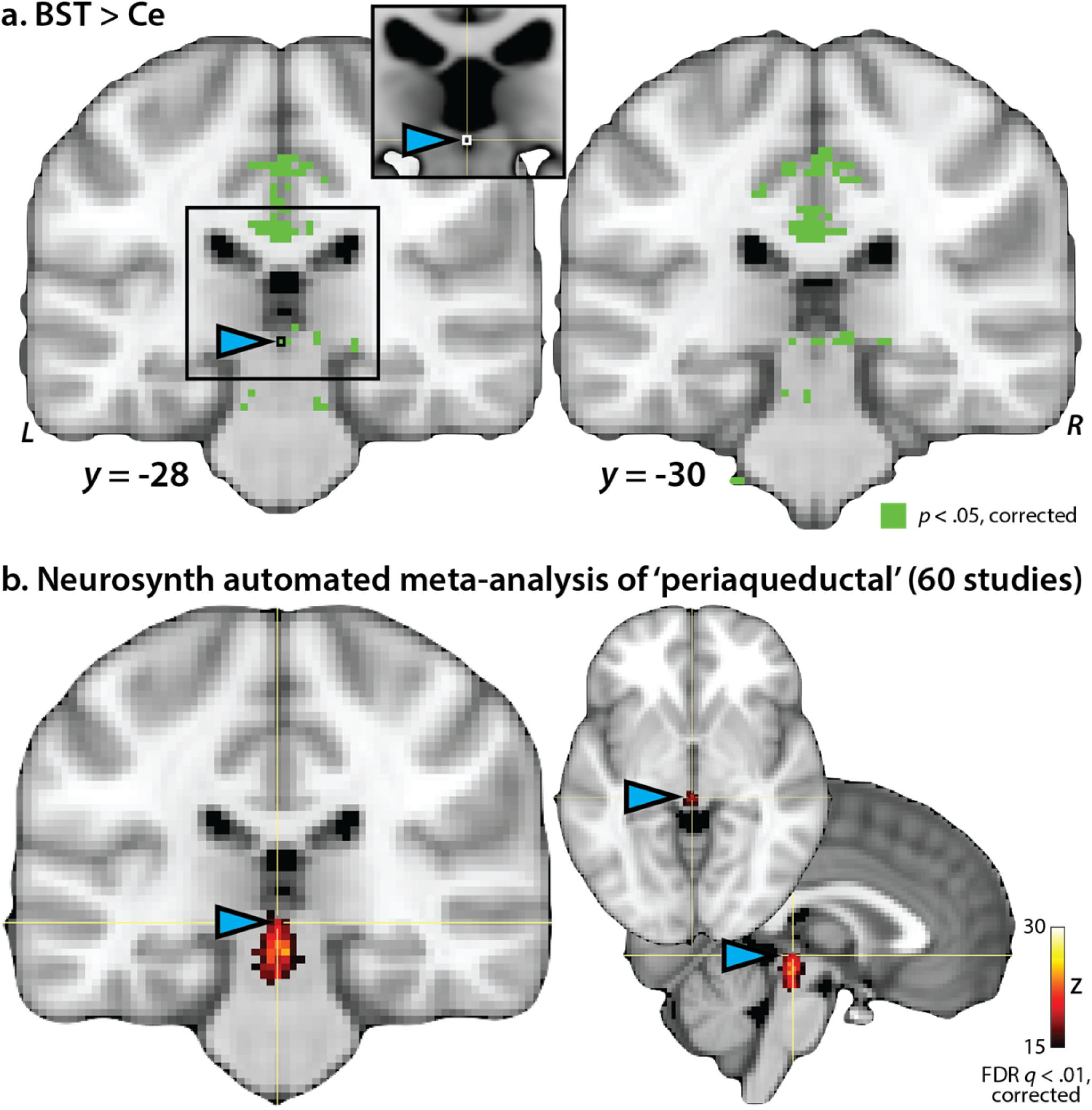
Relative to the Ce, the BST showed significantly greater coupling with a region of the brainstem in the region of the dorsal PAG. **a. BST vs. Ce contrast.** For maximal precision, this panel shows the uninterpolated, thresholded functional connectivity map displayed on the 2-mm MNI152 grid used for all analyses. The location of the peak voxel in the region of the dorsal BST is indicated by the *cyan* arrowhead (*x*=0, *y*=-28, *z*=-4, *t*=5.90, *p*<.05, corrected). This location lies within 1 mm of the PAG subdivisions recently identified by Ezra and colleagues using diffusion-weighted imaging (Ezra, Faull, Jbabdi, & Pattinson, 2015; their figure 3) and lies within the ‘full PAG’ mask of Coulombe and colleagues (Coulombe, Erpelding, Kucyi, & Davis, 2016). Inset depicts the corresponding location in the 1-mm MNI152 template. **b. Neurosynth automated meta-analysis of the term ‘periaqueductal’ (60 studies).** For illustrative purposes, this panel depicts the meta-analytic ‘forward inference’ map arbitrarily thresholded at approximately half the maximum value (*Z*>15, FDR *q*<.01, whole-brain corrected). Similar results have been previously reported using other meta-analytic approaches (Linnman, Moulton, Barmettler, Becerra, & Borsook, 2012). For example, using a manually curated database of 194 imaging studies, Linnman and colleagues reported that the mean (±SD) MNI coordinates for functional clusters labeled as PAG were *x*=|4| (±3), *y*=-29 (±5), *z*=-12 (±7). The location of the brainstem voxel highlighted in panel a is indicated by *cyan* arrowheads and the yellow cross-hair. Abbreviations—BST, bed nucleus of the stria terminalis; Ce, central nucleus of the amygdala; L, left hemisphere; PAG, periaqueductal gray; R, right hemisphere.

### Analyses Controlling for Regional Signal Quality

Functional connectivity is a complex metric that reflects the influence of both signal (i.e., the degree of regional coupling) and noise (Friston, 2011; Smith, 2012). To assess whether our results reflect variation in signal quality, we used a series of whole-brain regression analyses to estimate the functional connectivity of the BST and the Ce, as well as regional differences in connectivity, while co-varying for the quality of signal in the Ce and the BST seeds. Signal quality was estimated for each subject and seed using three widely used measures of functional data quality (e.g., Birn et al., 2014; Holmes et al., 2015): the temporal signal-to-noise ratio (tSNR; e.g., LaBar, Gitelman, Mesulam, & Parrish, 2001; Parrish, Gitelman, LaBar, & Mesulam, 2000), the amplitude of low frequency fluctuations (ALFF; square-root of the power in the 0.009-0.10 Hz pass-band; Zang et al., 2007; Zuo et al., 2010), and the fractional ALFF (fALFF; square-root of the power in the 0.009-0.10 Hz pass-band normalized by the total power across all frequencies; Zou et al., 2008; Zuo et al., 2010). Using a conventional analytic approach without additional nuisance variates (see **Table 5** in the main report), the BST showed significantly stronger coupling with the basal ganglia, thalamus, brainstem, and rostral cingulate extending into the vmPFC, whereas the Ce showed significantly stronger coupling with neighboring regions of the dorsal amygdala and anterior hippocampus (*p*<.05, corrected). This same pattern was evident for analyses that co-varied for mean-centered tSNR, ALFF, fALFF, and/or regional differences (e.g., BST_tSNR_-Ce_tSNR_), as illustrated in **Supplementary Figure S10**. These results indicate that the differences in intrinsic functional connectivity that we report (i.e., BST vs. Ce) are not driven by simple differences in regional signal quality. Nevertheless, as with any fMRI study focused on regional differences in connectivity—for example those focused on sub-divisions of the amygdala (e.g., Blackford et al., 2014; Etkin, Prater, Schatzberg, Menon, & Greicius, 2009; Gabard-Durnam et al., 2014; Qin et al., 2014; Qin, Young, Supekar, Uddin, & Menon, 2012; Roy et al., 2014; Roy et al., 2013) or of the central extended amygdala (Gorka, Torrisi, Shackman, Grillon, & Ernst, *in press*)—we cannot completely rule out the possibility that the differences in BST and Ce connectivity that we observed reflect more subtle differences in signal quality or reliability.

**Supplementary Figure S10.**
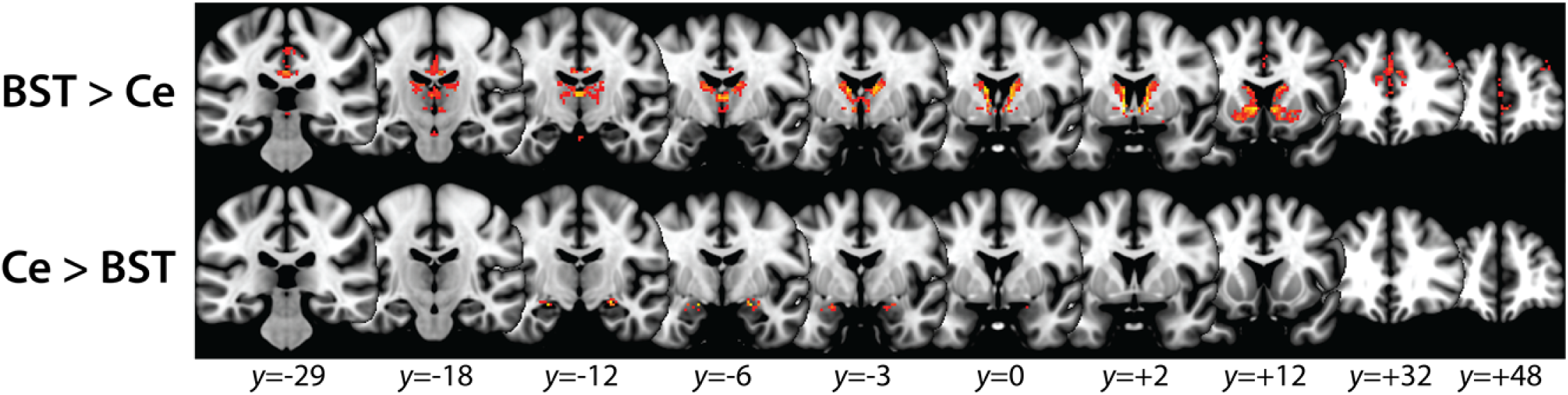
Differential functional connectivity of the BST vs. Ce controlling for ALFF and fALFF. Results of a paired *t*-test comparing the intrinsic functional connectivity of the BST and Ce controlling for nuisance variation in mean-centered BST_ALFF_, BST_fALFF_, Ce_ALFF_, and Ce_fALFF_. Conventions are similar to **Figure 4** in the main report, which depicts this same analysis without the additional covariates. Abbreviations—BST, bed nucleus of the stria terminalis; Ce, central nucleus of the amygdala; L, left hemisphere; R, right hemisphere.

## Footnotes

1 Specifically, for each subject, the de-faced T1 image was spatially normalized to the MNI152 template using the unified segmentation approach implemented in SPM12; (2) the 1-mm MNI152 template was de-faced to match the idiosyncratic de-facing of the T1 image; (3) the original T1 image was normalized to the individually de-faced 1-mm template using SyN; and (4) the inverse transformation was used to ‘reverse-normalize’ the MNI152 brain mask distributed with FSL to native space.

## REFERENCES

Acosta-Cabronero, J., Williams, G. B., Pereira, J. M., Pengas, G., & Nestor, P. J. (2008). The impact of skull-stripping and radio-frequency bias correction on grey-matter segmentation for voxel-based morphometry. Neuroimage, 39, 1654–1665.

Akam, T., & Kullmann, D. M. (2014). Oscillatory multiplexing of population codes for selective communication in the mammalian brain. Nature Reviews. Neuroscience, 15, 111–122.

Alheid, G. F., & Heimer, L. (1988). New perspectives in basal forebrain organization of special relevance for neuropsychiatric disorders: the striatopallidal, amygdaloid, and corticopetal components of substantia innominata. Neuroscience, 27, 1–39.

Alvarez, R. P., Chen, G., Bodurka, J., Kaplan, R., & Grillon, C. (2011). Phasic and sustained fear in humans elicits distinct patterns of brain activity. Neuroimage, 55, 389–400.

Alvarez, R. P., Kirlic, N., Misaki, M., Bodurka, J., Rhudy, J. L., Paulus, M. P., & Drevets, W. C. (2015). Increased anterior insula activity in anxious individuals is linked to diminished perceived control. Transl Psychiatry, 5, e591.

Amano, K., Tanikawa, T., Kawamura, H., Iseki, H., Notani, M., Kawabatake, H., … Kitamura, K. (1982). Endorphins and pain relief. Further observations on electrical stimulation of the lateral part of the periaqueductal gray matter during rostral mesencephalic reticulotomy for pain relief. Applied Neurophysiology, 45, 123–135.

Amunts, K., Kedo, O., Kindler, M., Pieperhoff, P., Mohlberg, H., Shah, N. J., … Zilles, K. (2005). Cytoarchitectonic mapping of the human amygdala, hippocampal region and entorhinal cortex: intersubject variability and probability maps. Anat Embryol, 210, 343–352.

Antoniadis, E. A., Winslow, J. T., Davis, M., & Amaral, D. G. (2007). Role of the primate amygdala in fear-potentiated startle: effects of chronic lesions in the rhesus monkey. Journal of Neuroscience, 27 (28), 7386–7396.

Ashburner, J., & Friston, K. J. (2005). Unified segmentation. Neuroimage, 26, 839–851.

Assareh, N., Sarrami, M., Carrive, P., & McNally, G. P. (2016). The organization of defensive behavior elicited by optogenetic excitation of rat lateral or ventrolateral periaqueductal gray. Behavioral Neuroscience, 130, 406–414.

Avants, B. B., Epstein, C. L., Grossman, M., & Gee, J. C. (2008). Symmetric diffeomorphic image registration with cross-correlation: Evaluating automated labeling of elderly and neurodegenerative brain. Medical Image Analysis, 12, 26–41.

Avants, B. B., Tustison, N. J., Song, G., Cook, P. A., Klein, A., & Gee, J. C. (2011). A reproducible evaluation of ANTs similarity metric performance in brain image registration. Neuroimage, 54, 2033–2044.

Avants, B. B., Yushkevich, P., Pluta, J., Minkoff, D., Korczykowski, M., Detre, J., & Gee, J. C. (2010). The optimal template effect in hippocampus studies of diseased populations. Neuroimage, 49, 2457–2466.

Avery, S. N., Clauss, J. A., & Blackford, J. U. (2016). The human BNST: Functional role in anxiety and addiction. Neuropsychopharmacology, 41, 126–141.

Avery, S. N., Clauss, J. A., Winder, D. G., Woodward, N., Heckers, S., & Blackford, J. U. (2014). BNST neurocircuitry in humans. Neuroimage, 91, 311–323.

Bandler, R., Price, J. L., & Keay, K. A. (2000). Brain mediation of active and passive emotional coping. Progress in Brain Research, 122, 333–349.

Banihashemi, L., Sheu, L. K., Midei, A. J., & Gianaros, P. J. (2015). Childhood physical abuse predicts stressor-evoked activity within central visceral control regions. Soc Cogn Affect Neurosci, 10, 474–485.

Bechara, A., Tranel, D., Damasio, H., Adolphs, R., Rockland, C., & Damasio, A. R. (1995). Double dissociation of conditioning and declarative knowledge relative to the amygdala and hippocampus in humans. Science, 269, 1115–1118.

Behzadi, Y., Restom, K., Liau, J., & Liu, T. T. (2007). A component based noise correction method (CompCor) for BOLD and perfusion based fMRI. Neuroimage, 37, 90–101.

Birn, R. M., Shackman, A. J., Oler, J. A., Williams, L. E., McFarlin, D. R., Rogers, G. M., … Kalin, N. H. (2014). Evolutionarily conserved dysfunction of prefrontal-amygdalar connectivity in early-life anxiety. Molecular Psychiatry, 19, 915–922.

Biswal, B., Yetkin, F. Z., Haughton, V. M., & Hyde, J. S. (1995). Functional connectivity in the motor cortex of resting human brain using echo-planar MRI. Magnetic Resonance in Medicine, 34, 537–541.

Brinkmann, L., Buff, C., Feldker, K., Neumeister, P., Heitmann, C. Y., Hofmann, D., … Straube, T. (under review/personal communication 7/20/2017). Inter-individual differences in trait anxiety shape the functional connectivity between the bed nucleus of the stria terminalis and the amygdala during brief threat processing. Neuroimage.

Brinkmann, L., Buff, C., Neumeister, P., Tupak, S. V., Becker, M. P., Herrmann, M. J., & Straube, T. (2017). Dissociation between amygdala and bed nucleus of the stria terminalis during threat anticipation in female post-traumatic stress disorder patients. Human Brain Mapping, 38, 2190–2205.

Buhle, J. T., Silvers, J. A., Wager, T. D., Lopez, R., Onyemekwu, C., Kober, H., … Ochsner, K. N. (2014). Cognitive reappraisal of emotion: A meta-analysis of human neuroimaging studies. Cerebral Cortex, 24, 2981–2990.

Button, K. S., Ioannidis, J. P., Mokrysz, C., Nosek, B. A., Flint, J., Robinson, E. S., & Munafo, M. R. (2013). Power failure: why small sample size undermines the reliability of neuroscience. Nature Reviews. Neuroscience, 14, 365–376.

Bystritsky, A. (2006). Treatment-resistant anxiety disorders. Molecular Psychiatry, 11, 805–814.

Cabral, J., Kringelbach, M. L., & Deco, G. (2014). Exploring the network dynamics underlying brain activity during rest. Progress in Neurobiology, 114C, 102–131.

Calhoon, G. G., & Tye, K. M. (2015). Resolving the neural circuits of anxiety. Nature Neuroscience, 18, 1394–1404.

Carrive, P., & Morgan, M. M. (2012). Periaqueductal gray. In J. K. Mai & G. Paxinos (Eds.), *The human nervous system*(3rd ed., pp. 367–400). New York: Academic Press.

Cavanagh, J. F., & Shackman, A. J. (2015). Frontal midline theta reflects anxiety and cognitive control: Meta-analytic evidence. Journal of Physiology, Paris, 109, 3–15.

Chang, L. J., Gianaros, P. J., Manuck, S. B., Krishnan, A., & Wager, T. D. (2015). A sensitive and specific neural signature for picture-induced negative affect. PLoS Biol, 13, e1002180.

Chen, S., Zhou, H., Guo, S., Zhang, J., Qu, Y., Feng, Z., … Zheng, X. (2015). Optogenetics based rat–robot control: Optical stimulation encodes “stop” and “escape” commands. Annals of Biomedical Engineering, 43, 1851–1864.

Cheng, D. T., Knight, D. C., Smith, C. N., & Helmstetter, F. J. (2006). Human amygdala activity during the expression of fear responses. Behavioral Neuroscience, 120, 1187–1195.

Cheng, D. T., Richards, J., & Helmstetter, F. J. (2007). Activity in the human amygdala corresponds to early, rather than late period autonomic responses to a signal for shock. Learning & Memory, 14, 485–490.

Choi, J. M., Padmala, S., & Pessoa, L. (2012). Impact of state anxiety on the interaction between threat monitoring and cognition. Neuroimage, 59, 1912–1923.

Choi, J. S., & Kim, J. J. (2010). Amygdala regulates risk of predation in rats foraging in a dynamic fear environment. Proceedings of the National Academy of Sciences of the United States of America, 107, 21773–21777.

Ciocchi, S., Herry, C., Grenier, F., Wolff, S. B., Letzkus, J. J., Vlachos, I., … Luthi, A. (2010). Encoding of conditioned fear in central amygdala inhibitory circuits. Nature, 468, 277–282.

Cloos, J. M., & Ferreira, V. (2009). Current use of benzodiazepines in anxiety disorders. Curr Opin Psychiatry, 22 (1), 90–95.

Costafreda, S. G., Brammer, M. J., David, A. S., & Fu, C. H. (2008). Predictors of amygdala activation during the processing of emotional stimuli: a meta-analysis of 385 PET and fMRI studies. Brain Research Reviews, 58, 57–70.

Coulombe, M. A., Erpelding, N., Kucyi, A., & Davis, K. D. (2016). Intrinsic functional connectivity of periaqueductal gray subregions in humans. Human Brain Mapping, 37, 1514–1530.

Cox, R. W. (1996). AFNI: Software for analysis and visualization of functional magnetic resonance neuroimages. Computers and Biomedical Research, 29, 162–173.

Davis, M., Walker, D. L., Miles, L., & Grillon, C. (2010). Phasic vs sustained fear in rats and humans: Role of the extended amygdala in fear vs anxiety. Neuropsychopharmacology, 35, 105–135.

Davis, M., & Whalen, P. J. (2001). The amygdala: vigilance and emotion. Molecular Psychiatry, 6, 13-34.

de la Vega, A., Chang, L. J., Banich, M. T., Wager, T. D., & Yarkoni, T. (2016). Large-scale meta-analysis of human medial frontal cortex reveals tripartite functional organization. Journal of Neuroscience, 36, 6553–6562.

deCampo, D. M., & Fudge, J. L. (2013). Amygdala projections to the lateral bed nucleus of the stria terminalis in the macaque: comparison with ventral striatal afferents. Journal of Comparative Neurology, 521, 3191–3216.

DiLuca, M., & Olesen, J. (2014). The cost of brain diseases: a burden or a challenge? Neuron, 82, 1205–1208.

Ehrlich, I., Humeau, Y., Grenier, F., Ciocchi, S., Herry, C., & Luthi, A. (2009). Amygdala inhibitory circuits and the control of fear memory. Neuron, 62, 757–771.

Entis, J. J., Doerga, P., Barrett, L. F., & Dickerson, B. C. (2012). A reliable protocol for the manual segmentation of the human amygdala and its subregions using ultra-high resolution MRI. Neuroimage, 60, 1226–1235.

Fadok, J. P., Krabbe, S., Markovic, M., Courtin, J., Xu, C., Massi, L., … Luthi, A. (2017). A competitive inhibitory circuit for selection of active and passive fear responses. Nature, 542, 96–100.

Faul, F., Erdfelder, E., Buchner, A., & Lang, A.-G. (2009). Statistical power analyses using G*Power 3.1: Tests for correlation and regression analyses. Behavior Research Methods, 41, 1149–1160.

Faul, F., Erdfelder, E., Lang, A.-G., & Buchner, A. (2007). G*Power 3: A flexible statistical power analysis program for the social, behavioral, and biomedical sciences. Behavior Research Methods, 39, 175–191.

Faull, O. K., & Pattinson, K. T. (2017). The cortical connectivity of the periaqueductal gray and the conditioned response to the threat of breathlessness. Elife, 6.

Fein, G., Landman, B., Tran, H., Barakos, J., Moon, K., Di Sclafani, V., & Shumway, R. (2006). Statistical parametric mapping of brain morphology: sensitivity is dramatically increased by using brain-extracted images as inputs. Neuroimage, 30, 1187–1195.

Feinstein, J. S., Adolphs, R., Damasio, A., & Tranel, D. (2011). The human amygdala and the induction and experience of fear. Current Biology, 21, 1–5.

Feinstein, J. S., Adolphs, R., & Tranel, D. (2016). A tale of survival from the world of Patient S.M. In D. G. Amaral & R. Adolphs (Eds.), Living without an amygdala. New York: Guilford.

Fischmeister, F. P., Hollinger, I., Klinger, N., Geissler, A., Wurnig, M. C., Matt, E., … Beisteiner, R. (2013). The benefits of skull stripping in the normalization of clinical fMRI data. Neuroimage Clin, 3, 369–380.

Fox, A. S., & Kalin, N. H. (2014). A translational neuroscience approach to understanding the development of social anxiety disorder and its pathophysiology. American Journal of Psychiatry, 171, 1162–1173.

Fox, A. S., Lapate, R. C., Davidson, R. J., & Shackman, A. J. (in press). Epilogue—The nature of emotion: A research agenda for the 21st century. In A. S. Fox, R. C. Lapate, A. J. Shackman & R. J. Davidson (Eds.), The nature of emotion. Fundamental questions (2nd ed., pp. [http://shackmanlab.org/wp-content/uploads/2017/2007/fox_shackman_NoE_Epilogue_070917Final.pdf]). New York: Oxford University Press.

Fox, A. S., Oler, J. A., Shackman, A. J., Shelton, S. E., Raveendran, M., McKay, D. R., … Kalin, N. H. (2015). Intergenerational neural mediators of early-life anxious temperament. Proceedings of the National Academy of Sciences USA, 112, 9118–9122.

Fox, A. S., Oler, J. A., Tromp, D. P., Fudge, J. L., & Kalin, N. H. (2015). Extending the amygdala in theories of threat processing. Trends in Neurosciences, 38, 319–329.

Fox, A. S., & Shackman, A. J. (*under review*). The central extended amygdala in fear and anxiety: Closing the gap between mechanistic and neuroimaging research [Invited Review]. Neuroscience Letters.

Fox, A. S., Shelton, S. E., Oakes, T. R., Converse, A. K., Davidson, R. J., & Kalin, N. H. (2010). Orbitofrontal cortex lesions alter anxiety-related activity in the primate bed nucleus of stria terminalis. Journal of Neuroscience, 30, 7023–7027.

Fox, A. S., Shelton, S. E., Oakes, T. R., Davidson, R. J., & Kalin, N. H. (2008). Trait-like brain activity during adolescence predicts anxious temperament in primates. PLoS ONE, 3, e2570.

Fox, M. D., Snyder, A. Z., Vincent, J. L., Corbetta, M., Van Essen, D. C., & Raichle, M. E. (2005). The human brain is intrinsically organized into dynamic, anticorrelated functional networks. Proceedings of the National Academy of Sciences of the United States of America, 102, 9673–9678.

Freese, J. L., & Amaral, D. G. (2009). Neuroanatomy of the primate amygdala. In P. J. Whalen & E. A. Phelps (Eds.), The human amygdala (pp. 3–42). NY: Guilford.

Fullana, M. A., Harrison, B. J., Soriano-Mas, C., Vervliet, B., Cardoner, N., Avila-Parcet, A., & Radua, J. (2016). Neural signatures of human fear conditioning: an updated and extended meta-analysis of fMRI studies. Molecular Psychiatry, 21, 500–508.

Fusar-Poli, P., Placentino, A., Carletti, F., Landi, P., Allen, P., Surguladze, S., … Politi, P. (2009). Functional atlas of emotional faces processing: a voxel-based meta-analysis of 105 functional magnetic resonance imaging studies. Journal of Psychiatry and Neuroscience, 34, 418–432.

Goode, T. D., & Maren, S. (2017). Role of the bed nucleus of the stria terminalis in aversive learning and memory. Learning and Memory, 24, 480–491.

Gorka, A. X., Torrisi, S., Shackman, A. J., Grillon, C., & Ernst, M. (*in press*). Intrinsic functional connectivity of the central nucleus of the amygdala and bed nucleus of the stria terminalis. Neuroimage.

Grayson, D. S., Bliss-Moreau, E., Machado, C. J., Bennett, J., Shen, K., Grant, K. A., … Amaral, D. G. (2016). The rhesus monkey connectome predicts disrupted functional networks resulting from pharmacogenetic inactivation of the amygdala. Neuron, 91, 453–466.

Greve, D. N., & Fischl, B. (2009). Accurate and robust brain image alignment using boundary-based registration. Neuroimage, 48, 63–72.

Griebel, G., & Holmes, A. (2013). 50 years of hurdles and hope in anxiolytic drug discovery. Nature Reviews. Drug Discovery, 12, 667–687.

Grupe, D. W., & Nitschke, J. B. (2013). Uncertainty and anticipation in anxiety: an integrated neurobiological and psychological perspective. Nature Reviews. Neuroscience, 14, 488–501.

Grupe, D. W., Oathes, D. J., & Nitschke, J. B. (2013). Dissecting the anticipation of aversion reveals dissociable neural networks. Cerebral Cortex, 23, 1874–1883.

Gungor, N. Z., & Paré, D. (2016). Functional heterogeneity in the bed nucleus of the stria terminalis. Journal of Neuroscience, 36, 8038–8049.

Hallquist, M. N., Hwang, K., & Luna, B. (2013). The nuisance of nuisance regression: spectral misspecification in a common approach to resting-state fMRI preprocessing reintroduces noise and obscures functional connectivity. Neuroimage, 82, 208–225.

Han, S., Soleiman, M. T., Soden, M. E., Zweifel, L. S., & Palmiter, R. D. (2015). Elucidating an affective pain circuit that creates a threat memory. Cell, 162, 363–374.

Herrmann, M. J., Boehme, S., Becker, M. P., Tupak, S. V., Guhn, A., Schmidt, B., … Straube, T. (2016). Phasic and sustained brain responses in the amygdala and the bed nucleus of the stria terminalis during threat anticipation. Human Brain Mapping, 37, 1091–1102.

Holmes, A. J., Hollinshead, M. O., O'Keefe, T. M., Petrov, V. I., Fariello, G. R., Wald, L. L., … Buckner, R. L.(2015). Brain Genomics Superstruct Project initial data release with structural, functional, and behavioral measures. Sci Data, 2, 150031.

Hrybouski, S., Aghamohammadi-Sereshki, A., Madan, C. R., Shafer, A. T., Baron, C. A., Seres, P., … Malykhin, N. V. (2016). Amygdala subnuclei response and connectivity during emotional processing. Neuroimage, 133, 98–110.

Iglesias, J. E., Liu, C. Y., Thompson, P., & Tu, Z. (2011). Robust brain extraction across datasets and comparison with publicly available methods. IEEE Transactions on Medical Imaging, 30, 1617–1634.

Izquierdo, A., Suda, R. K., & Murray, E. A. (2005). Comparison of the effects of bilateral orbital prefrontal cortex lesions and amygdala lesions on emotional responses in rhesus monkeys. Journal of Neuroscience, 25 (37), 8534–8542.

Jo, H. J., Gotts, S. J., Reynolds, R. C., Bandettini, P. A., Martin, A., Cox, R. W., & Saad, Z. S. (2013). Effective preprocessing procedures virtually eliminate distance-dependent motion artifacts in resting state FMRI. Journal of Applied Mathematics, 2013, 1–9.

Johnston, J. B. (1923). Further contributions to the study of the evolution of the forebrain. Journal of Comparative Neurology, 35, 337–481.

Kaczkurkin, A. N., Moore, T. M., Ruparel, K., Ciric, R., Calkins, M. E., Shinohara, R. T., … Satterthwaite, T. D. (2016). Elevated amygdala perfusion mediates developmental sex differences in trait anxiety. Biological Psychiatry, 80, 775–785.

Kalin, N. H. (2017). Mechanisms underlying the early risk to develop anxiety and depression: A translational approach. European Neuropsychopharmacology, 27, 543–553.

Kalin, N. H., Fox, A. S., Kovner, R., Riedel, M. K., Fekete, E. M., Roseboom, P. H., … Oler, J. A. (2016). Overexpressing corticotropin-releasing hormone in the primate amygdala increases anxious temperament and alters its neural circuit. Biological Psychiatry, 80, 345–355.

Kalin, N. H., Shelton, S. E., & Davidson, R. J. (2004). The role of the central nucleus of the amygdala in mediating fear and anxiety in the primate. Journal of Neuroscience, 24, 5506–5515.

Kalin, N. H., Shelton, S. E., & Davidson, R. J. (2007). Role of the primate orbitofrontal cortex in mediating anxious temperament. Biological Psychiatry, 62, 1134–1139.

Kalin, N. H., Shelton, S. E., Fox, A. S., Oakes, T. R., & Davidson, R. J. (2005). Brain regions associated with the expression and contextual regulation of anxiety in primates. Biological Psychiatry, 58, 796–804.

Kamali, A., Sair, H. I., Blitz, A. M., Riascos, R. F., Mirbagheri, S., Keser, Z., & Hasan, K. M. (2016). Revealing the ventral amygdalofugal pathway of the human limbic system using high spatial resolution diffusion tensor tractography. Brain Struct Funct, 221, 3561–3569.

Kamali, A., Yousem, D. M., Lin, D. D., Sair, H. I., Jasti, S. P., Keser, Z., … Hasan, K. M. (2015). Mapping the trajectory of the stria terminalis of the human limbic system using high spatial resolution diffusion tensor tractography. Neuroscience Letters, 608, 45–50.

Klein, A., Andersson, J., Ardekani, B. A., Ashburner, J., Avants, B., Chiang, M. C., … Parsey, R. V. (2009). Evaluation of 14 nonlinear deformation algorithms applied to human brain MRI registration. Neuroimage, 46, 786–802.

Klumpers, F., Kroes, M. C., Heitland, I., Everaerd, D., Akkermans, S. E., Oosting, R. S., … Baas, J. M. (2015). Dorsomedial prefrontal cortex mediates the impact of serotonin transporter linked polymorphic region genotype on anticipatory threat reactions. Biological Psychiatry, 78, 582–589.

Knight, D. C., Nguyen, H. T., & Bandettini, P. A. (2005). The role of the human amygdala in the production of conditioned fear responses. Neuroimage, 26, 1193–1200.

Korn, C. W., Vunder, J., Miró, J., Fuentemilla, L., Hurlemann, R., & Bach, D. R. (*in press*). Amygdala lesions reduce anxiety-like behavior in a human benzodiazepine-sensitive approach-avoidance conflict test. Biological Psychiatry.

Kragel, P. A., Knodt, A. R., Hariri, A. R., & LaBar, K. S. (2016). Decoding Spontaneous Emotional States in the Human Brain. PLoS Biol, 14, e2000106.

Kragel, P. A., & LaBar, K. S. (2015). Multivariate neural biomarkers of emotional states are categorically distinct. Soc Cogn Affect Neurosci, 10, 1437–1448.

LaBar, K. S., Gatenby, J. C., Gore, J. C., LeDoux, J. E., & Phelps, E. A. (1998). Human amygdala activation during conditioned fear acquisition and extinction: a mixed-trial fMRI study. Neuron, 20, 937–945.

Lange, M. D., Daldrup, T., Remmers, F., Szkudlarek, H. J., Lesting, J., Guggenhuber, S., … Pape, H. C. (*in press*). Cannabinoid CB1 receptors in distinct circuits of the extended amygdala determine fear responsiveness to unpredictable threat. Molecular Psychiatry.

LeDoux, J. E. (2000). Emotion circuits in the brain. Annual Review of Neuroscience, 23, 155–184.

LeDoux, J. E. (2007). The amygdala. Current Biology, 17, R868–874.

Li, H., Penzo, M. A., Taniguchi, H., Kopec, C. D., Huang, Z. J., & Li, B. (2013). Experience-dependent modification of a central amygdala fear circuit. Nature Neuroscience, 16, 332–339.

Lindquist, K. A., Satpute, A. B., Wager, T. D., Weber, J., & Barrett, L. F. (2016). The brain basis of positive and negative affect: Evidence from a meta-analysis of the human neuroimaging literature. Cerebral Cortex, 26, 1910–1922.

Logothetis, N. K. (2008). What we can do and what we cannot do with fMRI. Nature, 453, 869–878.

Mai, J. K., Paxinos, G., & Voss, T. (2007). Atlas of the human brain (3rd ed.). San Diego, CA: Academic Press.

Mason, W. A., Capitanio, J. P., Machado, C. J., Mendoza, S. P., & Amaral, D. G. (2006). Amygdalectomy and responsiveness to novelty in rhesus monkeys (Macaca mulatta): generality and individual consistency of effects. Emotion, 6, 73–81.

McMenamin, B. W., Langeslag, S. J., Sirbu, M., Padmala, S., & Pessoa, L. (2014). Network organization unfolds over time during periods of anxious anticipation. Journal of Neuroscience, 34, 11261–11273.

Mechias, M. L., Etkin, A., & Kalisch, R. (2010). A meta-analysis of instructed fear studies: implications for conscious appraisal of threat. Neuroimage, 49, 1760–1768.

Mobbs, D., Yu, R., Rowe, J. B., Eich, H., FeldmanHall, O., & Dalgleish, T. (2010). Neural activity associated with monitoring the oscillating threat value of a tarantula. Proceedings of the National Acadademy of Sciences USA, 107, 20582–20586.

Motta, S. C., Carobrez, A. P., & Canteras, N. S. (2017). The periaqueductal gray and primal emotional processing critical to influence complex defensive responses, fear learning and reward seeking. Neuroscience and Biobehavioral Reviews, 76, 39–47.

Motzkin, J. C., Philippi, C. L., Oler, J. A., Kalin, N. H., Baskaya, M. K., & Koenigs, M. (2015). Ventromedial prefrontal cortex damage alters resting blood flow to the bed nucleus of stria terminalis. Cortex, 64, 281–288.

Nacewicz, B. M., Alexander, A. L., Kalin, N. H., & Davidson, R. J. (2014). The neurochemical underpinnings of human amygdala volume including subregional contributions. Biological Psychiatry, 75, S222.

Najafi, M., Kinnison, J., & Pessoa, L. (2017). Intersubject brain network organization during dynamic anxious anticipation. bioRxiv.

Nashold, B. S., Wilson, W. P., & Slaughter, D. G. (1969). Sensations evoked by stimulation in the midbrain of man. Journal of Neurosurgery, 30, 14–24.

Nauta, W. J. (1961). Fibre degeneration following lesions of the amygdaloid complex in the monkey. Journal of Anatomy, 95, 515–531.

Nichols, T., Brett, M., Andersson, J., Wager, T., & Poline, J. B. (2005). Valid conjunction inference with the minimum statistic. Neuroimage, 25(3), 653–660.

Nooner, K. B., Colcombe, S. J., Tobe, R. H., Mennes, M., Benedict, M. M., Moreno, A. L., … Milham, M. P. (2012). The NKI-Rockland sample: A model for accelerating the pace of discovery science in psychiatry. Front Neurosci, 6, 152.

Nummenmaa, L., & Saarimaki, H. (in press). Emotions as discrete patterns of systemic activity. Neuroscience Letters

Oler, J. A., Birn, R. M., Patriat, R., Fox, A. S., Shelton, S. E., Burghy, C. A., … Kalin, N. H. (2012). Evidence for coordinated functional activity within the extended amygdala of non-human and human primates. Neuroimage, 61, 1059–1066.

Oler, J. A., Fox, A. S., Shackman, A. J., & Kalin, N. H. (2016). The central nucleus of the amygdala is a critical substrate for individual differences in anxiety. In D. G. Amaral & R. Adolphs (Eds.), Living without an amygdala. NY: Guilford.

Oler, J. A., Fox, A. S., Shelton, S. E., Rogers, J., Dyer, T. D., Davidson, R. J., … Kalin, N. H. (2010). Amygdalar and hippocampal substrates of anxious temperament differ in their heritability. Nature, 466, 864–868.

Oler, J. A., Tromp, D. P., Fox, A. S., Kovner, R., Davidson, R. J., Alexander, A. L., … Fudge, J. L. (2017). Connectivity between the central nucleus of the amygdala and the bed nucleus of the stria terminalis in the non-human primate: neuronal tract tracing and developmental neuroimaging studies. Brain Struct Funct, 222, 21–39.

Ongur, D., & Price, J. L. (2000). The organization of networks within the orbital and medial prefrontal cortex of rats, monkeys and humans. Cerebral Cortex, 10, 206–219.

Pare, D., & Duvarci, S. (2012). Amygdala microcircuits mediating fear expression and extinction. Current Opinion in Neurobiology, 22, 717–723.

Pedersen, W. S., Balderston, N. L., Miskovich, T. A., Belleau, E. L., Helmstetter, F. J., & Larson, C. L. (2017). The effects of stimulus novelty and negativity on BOLD activity in the amygdala, hippocampus, and bed nucleus of the stria terminalis. Soc Cogn Affect Neurosci, 12, 748–757.

Penzo, M. A., Robert, V., & Li, B. (2014). Fear conditioning potentiates synaptic transmission onto long-range projection neurons in the lateral subdivision of central amygdala. Journal of Neuroscience, 34, 2432–2437.

Penzo, M. A., Robert, V., Tucciarone, J., De Bundel, D., Wang, M., Van Aelst, L., … Li, B. (2015). The paraventricular thalamus controls a central amygdala fear circuit. Nature, 519, 455–459.

Pessoa, L. (2017). A network model of the emotional brain. Trends in Cognitive Sciences, 21, 357–371.

Poldrack, R. A., Baker, C. I., Durnez, J., Gorgolewski, K. J., Matthews, P. M., Munafo, M. R., … Yarkoni, T. (2017). Scanning the horizon: towards transparent and reproducible neuroimaging research. Nature Reviews. Neuroscience, 18, 115–126.

Power, J. D., Schlaggar, B. L., & Petersen, S. E. (2015). Recent progress and outstanding issues in motion correction in resting state fMRI. Neuroimage, 105, 536–551.

Richardson, D. E., & Akil, H. (1977). Pain reduction by electrical brain stimulation inman. Part 1: Acute administration in periaqueductal and periventricular sites. Journal of Neurosurgery, 47, 178–183.

Roesch, M. R., Esber, G. R., Li, J., Daw, N. D., & Schoenbaum, G. (2012). Surprise! Neural correlates of Pearce-Hall and Rescorla-Wagner coexist within the brain. European Journal of Neuroscience, 35, 1190–1200.

Rudebeck, P. H., Saunders, R. C., Prescott, A. T., Chau, L. S., & Murray, E. A. (2013). Prefrontal mechanisms of behavioral flexibility, emotion regulation and value updating. Nature Neuroscience, 16, 1140–1145.

Sabatinelli, D., Fortune, E. E., Li, Q., Siddiqui, A., Krafft, C., Oliver, W. T., … Jeffries, J. (2011). Emotional perception: Meta-analyses of face and natural scene processing. Neuroimage, 54, 2524–2533.

Salomon, J. A., Haagsma, J. A., Davis, A., de Noordhout, C. M., Polinder, S., Havelaar, A. H., … Vos, T. (2015). Disability weights for the Global Burden of Disease 2013 study. Lancet Glob Health, 3, e712–723.

Sato, M., Ito, M., Nagase, M., Sugimura, Y. K., Takahashi, Y., Watabe, A. M., & Kato, F. (2015). The lateral parabrachial nucleus is actively involved in the acquisition of fear memory in mice. Mol Brain, 8, 22.

Satpute, A. B., Wager, T. D., Cohen-Adad, J., Bianciardi, M., Choi, J. K., Buhle, J. T., … Barrett, L. F. (2013). Identification of discrete functional subregions of the human periaqueductal gray. Proceedings of the National Academy of Sciences of the United States of America, 110, 17101–17106.

Sergerie, K., Chochol, C., & Armony, J. L. (2008). The role of the amygdala in emotional processing: a quantitative meta-analysis of functional neuroimaging studies. Neuroscience and Biobehavioral Reviews, 32, 811–830.

Shackman, A. J., & Fox, A. S. (2016). Contributions of the central extended amygdala to fear and anxiety. Journal of Neuroscience, 36, 8050–8063.

Shackman, A. J., & Fox, A. S. (*in press*). Afterword: How are emotions organized in the brain? In A. S. Fox, R. C. Lapate, A. J. Shackman & R. J. Davidson (Eds.), The nature of emotion. Fundamental questions (2nd ed.). New York, NY: Oxford University Press.

Shackman, A. J., Fox, A. S., Oler, J. A., Shelton, S. E., Davidson, R. J., & Kalin, N. H. (2013). Neural mechanisms underlying heterogeneity in the presentation of anxious temperament. Proceedings of the National Academy of Sciences of the United States of America, 110, 6145–6150.

Shackman, A. J., Fox, A. S., Oler, J. A., Shelton, S. E., Oakes, T. R., Davidson, R. J., & Kalin, N. H. (2017). Heightened extended amygdala metabolism following threat characterizes the early phenotypic risk to develop anxiety-related psychopathology. Molecular Psychiatry, 22, 724–732.

Shackman, A. J., Fox, A. S., & Seminowicz, D. A. (2015). The cognitive-emotional brain: Opportunities and challenges for understanding neuropsychiatric disorders. Behavioral and Brain Sciences, 38, e86.

Shackman, A. J., Kaplan, C. M., Stockbridge, M. D., Tillman, R. M., Tromp, D. P. M., Fox, A. S., & Gamer, M.(2016). The neurobiology of anxiety and attentional biases to threat: Implications for understanding anxiety disorders in adults and youth. Journal of Experimental Psychopathology, 7, 311–342.

Shackman, A. J., McMenamin, B. W., Maxwell, J. S., Greischar, L. L., & Davidson, R. J. (2009). Right dorsolateral prefrontal cortical activity and behavioral inhibition. Psychological Science, 20, 1500–1506.

Shackman, A. J., Salomons, T. V., Slagter, H. A., Fox, A. S., Winter, J. J., & Davidson, R. J. (2011). The integration of negative affect, pain and cognitive control in the cingulate cortex. Nature Reviews. Neuroscience, 12, 154–167.

Shackman, A. J., Tromp, D. P. M., Stockbridge, M. D., Kaplan, C. M., Tillman, R. M., & Fox, A. S. (2016). Dispositional negativity: An integrative psychological and neurobiological perspective. Psychological Bulletin, 142, 1275–1314.

Shattuck, D. W., Sandor-Leahy, S. R., Schaper, K. A., Rottenberg, D. A., & Leahy, R. M. (2001). Magnetic resonance image tissue classification using a partial volume model. Neuroimage, 13, 856–876.

Šidák, Z. K. (1967). Rectangular confidence regions for the means of multivariate normal distributions. Journal of the American Statistical Association, 62, 626–633.

Siegel, J. S., Power, J. D., Dubis, J. W., Vogel, A. C., Church, J. A., Schlaggar, B. L., & Petersen, S. E. (2014). Statistical improvements in functional magnetic resonance imaging analyses produced by censoring high-motion data points. Human Brain Mapping, 35, 1981–1996.

Sladky, R., Geissberger, N., Pfabigan, D. M., Kraus, C., Tik, M., Woletz, M., … Windischberger, C. (*in press*). Unsmoothed functional MRI of the human amygdala and bed nucleus of the stria terminalis during processing of emotional faces. Neuroimage.

Smith, S. M. (2002). Fast robust automated brain extraction. Human Brain Mapping, 17, 143–155.

Somerville, L. H., Wagner, D. D., Wig, G. S., Moran, J. M., Whalen, P. J., & Kelley, W. M. (2013). Interactions between transient and sustained neural signals support the generation and regulation of anxious emotion. Cerebral Cortex, 23, 49–60.

Somerville, L. H., Whalen, P. J., & Kelley, W. M. (2010). Human bed nucleus of the stria terminalis indexes hypervigilant threat monitoring. Biological Psychiatry, 68, 416–424.

Stelzer, J., Lohmann, G., Mueller, K., Buschmann, T., & Turner, R. (2014). Deficient approaches to human neuroimaging. Front Hum Neurosci, 8, 462.

Stevens, J. S., Kim, Y. J., Galatzer-Levy, I. R., Reddy, R., Ely, T. D., Nemeroff, C. B., … Ressler, K. J. (2017). Amygdala reactivity and anterior cingulate habituation predict posttraumatic stress disorder symptom maintenance after acute civilian trauma. Biological Psychiatry, 81, 1023–1029.

Stocker, T. (2007). On the asymptotic bias of OLS in dynamic regression models with autocorrelated errors. Statistical Papers, 48, 81–93.

Stout, D. M., Shackman, A. J., Pedersen, W. S., Miskovich, T. A., & Larson, C. L. (*in press*). Neural circuitry governing anxious individuals' mis-allocation of working memory to threat. Scientific Reports.

Theiss, J. D., Ridgewell, C., McHugo, M., Heckers, S., & Blackford, J. U. (2017). Manual segmentation of the human bed nucleus of the stria terminalis using 3T MRI. Neuroimage, 146, 288–292.

Torrisi, S., O'Connell, K., Davis, A., Reynolds, R., Balderston, N., Fudge, J. L., … Ernst, M. (2015). Resting state connectivity of the bed nucleus of the stria terminalis at ultra-high field. Human Brain Mapping, 36, 4076–4088.

Tovote, P., Esposito, M. S., Botta, P., Chaudun, F., Fadok, J. P., Markovic, M., … Luthi, A. (2016). Midbrain circuits for defensive behaviour. Nature, 534, 206–212.

Tovote, P., Fadok, J. P., & Luthi, A. (2015). Neuronal circuits for fear and anxiety. Nature Reviews. Neuroscience, 16, 317–331.

Turner, R., & Geyer, S. (2014). Comparing like with like: the power of knowing where you are. Brain Connect, 4, 547–557.

Tustison, N. J., Cook, P. A., Klein, A., Song, G., Das, S. R., Duda, J. T., … Avants, B. B. (2014). Large-scale evaluation of ANTs and FreeSurfer cortical thickness measurements. Neuroimage, 99, 166–179.

Tyszka, J. M., & Pauli, W. M. (2016). In vivo delineation of subdivisions of the human amygdaloid complex in a high-resolution group template. Human Brain Mapping, 37, 3979–3998.

Uddin, L. Q., Kinnison, J., Pessoa, L., & Anderson, M. L. (2014). Beyond the tripartite cognition-emotion-interoception model of the human insular cortex. Journal of Cognitive Neuroscience, 26, 16–27.

van Well, S., Visser, R. M., Scholte, H. S., & Kindt, M. (2012). Neural substrates of individual differences in human fear learning: evidence from concurrent fMRI, fear-potentiated startle, and US-expectancy data. Cogn Affect Behav Neurosci, 12, 499–512.

Wager, T. D., Kang, J., Johnson, T. D., Nichols, T. E., Satpute, A. B., & Barrett, L. F. (2015). A Bayesian model of category-specific emotional brain responses. PLoS Comput Biol, 11, e1004066.

Wang, K., Gaitsch, H., Poon, H., Cox, N. J., & Rzhetsky, A. (in press). Classification of common human diseases derived from shared genetic and environmental determinants. Nature Genetics.

Whiteford, H. A., Degenhardt, L., Rehm, J., Baxter, A. J., Ferrari, A. J., Erskine, H. E., … Vos, T. (2013). Global burden of disease attributable to mental and substance use disorders: findings from the Global Burden of Disease Study 2010. Lancet, 382, 1575–1586.

Wiegert, J. S., Mahn, M., Prigge, M., Printz, Y., & Yizhar, O. (2017). Silencing neurons: Tools, applications, and experimental constraints. Neuron, 95, 504–529.

Wise, R. A., & Koob, G. F. (2014). The development and maintenance of drug addiction. Neuropsychopharmacology, 39 (2), 254–262.

Wood, K. H., Ver Hoef, L. W., & Knight, D. C. (2014). The amygdala mediates the emotional modulation of threat-elicited skin conductance response. Emotion, 14, 693–700.

Yilmazer-Hanke, D. M. Amygdala. (2012). In J. K. Mai & G. Paxinos (Eds.), The human nervous system (pp. 759–834). San Diego: Academic Press.

Yu, K., Ahrens, S., Zhang, X., Schiff, H., Ramakrishnan, C., Fenno, L., … Li, B. (2017). The central amygdala controls learning in the lateral amygdala. bioRxiv.

## REFERENCES

Fox, A. S., Lapate, R. C., Davidson, R. J., & Shackman, A. J. (*in press*). Epilogue—The nature of emotion: A research agenda for the 21st century. In A. S. Fox, R. C. Lapate, A. J. Shackman & R. J. Davidson (Eds.), The nature of emotion. Fundamental questions (2nd ed., pp. [http://shackmanlab.org/wp-content/uploads/2017/2007/fox_shackman_NoE_Epilogue_070917Final.pdf]). New York: Oxford University Press.

Gorka, A. X., Torrisi, S., Shackman, A. J., Grillon, C., & Ernst, M. (in press). Intrinsic functional connectivity of the central nucleus of the amygdala and bed nucleus of the stria terminalis. Neuroimage.

Ongur, D., Ferry, A. T., & Price, J. L. (2003). Architectonic subdivision of the human orbital and medial prefrontal cortex. Journal of Comparative Neurology, 460, 425–449.

## SUPPLEMENTARY REFERENCES

Ashburner, J., & Friston, K. J. (2005). Unified segmentation. Neuroimage, 26, 839–851.

Blackford, J. U., Clauss, J. A., Avery, S. N., Cowan, R. L., Benningfield, M. M., & VanDerKlok, R. M. (2014). Amygdala-cingulate intrinsic connectivity is associated with degree of social inhibition. Biological Psychology, 99, 15–25.

Cheng, W., Dryden, I. L., & Huang, X. (2016). Bayesian registration of functions and curves. Bayesian Analysis, 11, 447–475.

Chung, M. K., Worsley, K. J., Nacewicz, B. M., Dalton, K. D., & Davidson, R. J. (2010). General multivariate linear modeling of surface shapes using SurfStat. Neuroimage, 53, 491–505.

Etkin, A., Prater, K. E., Schatzberg, A. F., Menon, V., & Greicius, M. D. (2009). Disrupted amygdalar subregion functional connectivity and evidence of a compensatory network in generalized anxiety disorder. Archives of General Psychiatry, 66, 1361–1372.

Ezra, M., Faull, O. K., Jbabdi, S., & Pattinson, K. T. (2015). Connectivity-based segmentation of the periaqueductal gray matter in human with brainstem optimized diffusion MRI. Human Brain Mapping, 36, 3459–3471.

Friston, K. J. (2011). Functional and effective connectivity: A review. Brain Connectivity, 1, 13–36.

Gabard-Durnam, L. J., Flannery, J., Goff, B., Gee, D. G., Humphreys, K. L., Telzer, E., … Tottenham, N. (2014). The development of human amygdala functional connectivity at rest from 4 to 23 years: a cross-sectional study. Neuroimage, 95, 193–207.

Holmes, A. J., Hollinshead, M. O., O'Keefe, T. M., Petrov, V. I., Fariello, G. R., Wald, L. L., … Buckner, R. L. (2015). Brain Genomics Superstruct Project initial data release with structural, functional, and behavioral measures. Sci Data, 2, 150031.

Kruger, O., Shiozawa, T., Kreifelts, B., Scheffler, K., & Ethofer, T. (2015). Three distinct fiber pathways of the bed nucleus of the stria terminalis to the amygdala and prefrontal cortex. Cortex, 66, 60–68.

Kurtek, S. (2017). A geometric approach to pairwise Bayesian alignment of functional data using importance sampling. Electron. J. Statist., 11, 502-531.

LaBar, K. S., Gitelman, D. R., Mesulam, M. M., & Parrish, T. B. (2001). Impact of signal-to-noise on functional MRI of the human amygdala. Neuroreport, 12, 3461–3464.

Linnman, C., Moulton, E. A., Barmettler, G., Becerra, L., & Borsook, D. (2012). Neuroimaging of the periaqueductal gray: state of the field. Neuroimage, 60, 505–522.

Nacewicz, B. M., Dalton, K. M., Johnstone, T., Long, M. T., McAuliff, E. M., Oakes, T. R., … Davidson, R. J.(2006). Amygdala volume and nonverbal social impairment in adolescent and adult males with autism. Archives of General Psychiatry, 63, 1417–1428.

Parrish, T. B., Gitelman, D. R., LaBar, K. S., & Mesulam, M. M. (2000). Impact of signal-to-noise on functional MRI. Magnetic Resonance in Medicine, 44, 925–932.

Prevost, C., McCabe, J. A., Jessup, R. K., Bossaerts, P., & O'Doherty, J. P. (2011). Differentiable contributions of human amygdalar subregions in the computations underlying reward and avoidance learning. European Journal of Neuroscience, 34, 134–145.

Qin, S., Young, C. B., Duan, X., Chen, T., Supekar, K., & Menon, V. (2014). Amygdala subregional structure and intrinsic functional connectivity predicts individual differences in anxiety during early childhood. Biological Psychiatry, 75(11), 892–900.

Qin, S., Young, C. B., Supekar, K., Uddin, L. Q., & Menon, V. (2012). Immature integration and segregation of emotion-related brain circuitry in young children. Proceedings of the National Academy of Sciences of the United States of America, 109, 7941–7946.

Roy, A. K., Benson, B. E., Degnan, K. A., Perez-Edgar, K., Pine, D. S., Fox, N. A., & Ernst, M. (2014). Alterations in amygdala functional connectivity reflect early temperament. Biological Psychology, 103, 248–254.

Roy, A. K., Fudge, J. L., Kelly, C., Perry, J. S., Daniele, T., Carlisi, C., … Ernst, M. (2013). Intrinsic functional connectivity of amygdala-based networks in adolescent generalized anxiety disorder. Journal of the American Academy of Child and Adolescent Psychiatry, 52, 290–299 e292.

Smith, S. M. (2002). Fast robust automated brain extraction. Human Brain Mapping, 17, 143–155.

Smith, S. M. (2012). The future of FMRI connectivity. Neuroimage, 62, 1257–1266.

Tucker, J. D., Wu, W., & Srivastava, A. (2013). Generative models for functional data using phase and amplitude separation. Computational Statistics & Data Analysis, 61, 50–66.

Wu, W., & Srivastava, A. (2014). Analysis of spike train data: Alignment and comparisons using the extended Fisher-Rao metric. Electronic Journal of Statistics, 8, 1776–1785.

Yarkoni, T., Poldrack, R. A., Nichols, T. E., Van Essen, D. C., & Wager, T. D. (2011). Large-scale automated synthesis of human functional neuroimaging data. Nat Methods, 8, 665–670.

Zang, Y. F., He, Y., Zhu, C. Z., Cao, Q. J., Sui, M. Q., Liang, M., … Wang, Y. F. (2007). Altered baseline brain activity in children with ADHD revealed by resting-state functional MRI. Brain and Development, 29, 83–91.

Zou, Q. H., Zhu, C. Z., Yang, Y., Zuo, X. N., Long, X. Y., Cao, Q. J., … Zang, Y. F. (2008). An improved approach to detection of amplitude of low-frequency fluctuation (ALFF) for resting-state fMRI: fractional ALFF. Journal of Neuroscience Methods, 172, 137–141.

Zuo, X. N., Di Martino, A., Kelly, C., Shehzad, Z. E., Gee, D. G., Klein, D. F., … Milham, M. P. (2010). The oscillating brain: complex and reliable. Neuroimage, 49, 1432–1445.

